# Aging-enhanced accumulation of fibroblasts excludes oligodendrocytes in demyelinated lesions

**DOI:** 10.1101/2025.07.27.667059

**Authors:** Brian M. Lozinski, George S. Melchor, Charlotte D’Mello, Rianne P. Gorter, Yifei Dong, Samira Ghorbani, Cenxiao Li, Marlene Morch, Dorsa Moezzi, Rajiv Jain, Parisa Etemadi Nezhad, Chang-Chun Ling, Carlos Camara-Lemarroy, Rui Li, Jeffrey K. Huang, V. Wee Yong

## Abstract

Fibroblast dysregulation contributes to pathological fibrosis and aberrant repair. Emerging evidence suggest that fibroblasts accumulate in lesions following central nervous system injury, but whether and how they influence oligodendrocyte repair responses, including in aging, is uncertain. Here we report that fibroblasts accumulate in the parenchyma of spinal cord white matter lesions of 6 – 10 week old young mice after lysolecithin-induced demyelination. This was first observed through immunofluorescence microscopy that employed several markers attributed to fibroblasts, including platelet-derived growth factor-β, collagen type 1α1, α-smooth muscle actin, periostin and fibronectin; and by the use of platelet-derived growth factor-β TdTomato reporter transgenic mice. Single-nucleus and spatial transcriptomics of lysolecithin lesions established the presence of fibroblasts in lysolecithin lesions and delineated them from closely related pericytes. CellChat ligand – receptor analyses highlight fibroblasts in the lysolecithin environment as a major source of input of signals for microglia/macrophages and oligodendrocyte precursor cells, with numerous reciprocal interactions. The infiltration of fibroblasts was promoted by microglia/macrophages, as anticipated by their temporal representation in lysolecithin lesions, and by tissue culture experiments where the migration of fibroblasts was enhanced by macrophages. Of particular relevance to spontaneous regenerative events in lysolecithin demyelination, areas of fibroblast accumulation were devoid of oligodendrocyte precursor cells. In tissue culture, oligodendrocyte precursor cells were excluded from fibroblast domains. Moreover, fibroblast accumulation after lysolecithin injury was enhanced with increasing age, a known detriment to the capacity to remyelinate after injury, and exclusion of oligodendrocyte precursor cells from fibroblast areas of 48 - 52 week mice exceeds that occurring in younger 6 - 10 weeks animals. Finally, by mining a publicly available single-nucleus RNA database of multiple sclerosis, we found fibroblasts in the edge of chronic active and chronic inactive lesions and in lesion core, and fewer in periplaque or normal white matter. We identified several communication networks between fibroblasts, microglia/macrophages and oligodendrocyte precursor cells in these MS lesions. Our collective results demonstrate a role of fibroblasts in demyelination-associated neuropathology, which is exacerbated by aging, and highlight the importance of regulating fibroblasts to promote effective CNS repair.

## Introduction

Fibroblasts are a heterogeneous cell population important for synthesis of extracellular matrix (ECM) components, maintenance of cellular niches, tissue regeneration and wound healing^1^. In these roles, fibroblasts communicate with immune cells such as macrophages. Fibroblasts associated with the CNS primarily reside in the meninges, perivascular space, and choroid plexus where they contribute to tissue homeostasis^2^ and immune regulation^3,4^. Several recent studies suggest that in response to neural injury, these border-associated fibroblasts accumulate in the CNS parenchyma and contribute to chronic fibrotic scaring^3,5–11^. Nonetheless, beneficial roles of fibroblasts may manifest in the acute phase after neural injury, and include regenerative functions and limiting the expansion of lesions^9,12^. Overall, while fibroblast activation is necessary to maintain tissue integrity, it may also ultimately impair regenerative processes^3,13,14^.

Remyelination is the most common and robust form of CNS regeneration^15–17^, involving the migration, proliferation, and differentiation of oligodendrocyte progenitor cells (OPCs) into mature myelinating oligodendrocytes^18,19^. Restoring myelin to denuded axons reconstitutes saltatory conduction, axonal trophic support, and produces a mechanical barrier to injury leading to axonal preservation and functional improvement^15,20,21^. However, remyelination often fails due to extrinsic variables in the environment, such as inhibitory ECM, and intrinsic factors including age-related cellular dysfunctions^21–23^.

Age contributes to disease worsening across a range of conditions, including MS and fibrotic disorders^24–26^. OPCs also show signs of age-related impairment, including upregulation of senescence markers, epigenetic dysfunction, reduced responsiveness to growth factors, and impaired differentiation and remyelination capacity^23,27–29^. While age-related decline in remyelination is well recognized, the impact of aging on fibroblast responses to CNS injury remains poorly understood.

In this study, we characterized the response of fibroblasts in lesions following focal lysolecithin (LPC)-mediated demyelination of the spinal cord white matter, and further evaluated the influence of increasing age on the fibrotic response. We compared middle-aged to young mice, as middle-age is a period when many people with MS transition to progressive MS. With single-nucleus and spatial transcriptomics of LPC tissue and cells in culture, we interrogated interactions between fibroblasts, microglia/macrophages and OPCs. Finally, we utilized a publicly available transcriptomic dataset to show that fibroblasts are present in MS lesions, and describe communication patterns between fibroblasts, microglia/macrophages and OPCs in MS. Our findings highlight the necessity of considering invasion of fibroblasts into the CNS parenchyma in lesion evolution and recovery.

## Materials and methods

### MS specimens

Postmortem frozen brain tissues from people with MS were obtained from The Montreal Brain Bank courtesy of Dr. Alexandre Prat, with informed consent approved by the CRCHUM and University of Montreal research ethics committee. Additional autopsied frozen MS brain tissues were from The Multiple Sclerosis and Parkinson’s Tissue Bank situated at Imperial College, London. The use of these human tissue in Calgary for research was approved by the Conjoint Health Research Ethics Board at the University of Calgary (Ethics ID REB15-0444). MS sections were characterized using luxol fast blue (LFB) and hematoxylin & eosin (H&E) to identify lesions as previously described^30^.

### Mice

All experiments were conducted with ethics approval from the Animal Care Committee at the University of Calgary under regulations of the Canadian Council of Animal Care. Female C57BL/6J mice were from Jackson laboratories. CX3CR1-CreER (JAX 021160), PDGFRβ-P2A-CreERT2 (JAX 030201), Ai9 [Rosa26-TdTomato (JAX 007905)], NG2-CreER (JAX 008538), and MAPT(Tau)-mGFP (JAX 021162) mice were from The Jackson Laboratory. We crossed PDGFRβCreER with Ai9 mice, where progenies after tamoxifen induction were referred as PDGFRβ^TdTom^ mice. NG2-CreER and MAPT-mGFP mice were bred to produce NG2^CreER^:MAPT^mGFP^ mice. All mouse strains were bred in the single barrier mouse unit at the University of Calgary. All young female mice were 6-10 weeks of age, and all middle-aged female mice were 48-52 weeks of age. Male and female CD1 pups P0-2 were used for OPC and meningeal fibroblast cultures. C57BL6 female mice 6-10 weeks of age were used for bone marrow-derived macrophages (BMDM) cultures. All mice were maintained on a 12-h light/dark cycle with food (Pico-Vac Mouse Diet 20) and water given ad libitum.

### Spinal cord surgery

Lysolecithin/lysophosphatidylcholine (LPC) demyelination was accomplished as previously described^31^. Mice were anaesthetized with intraperitoneal injections of ketamine (100 mgkg-1) and xylazine (10 mgkg-1). Buprenorphine (0.05 mgkg-1) was given as an analgesic. Mice were positioned on a stereotaxic frame and a 2-3 cm incision was made between the shoulder blades. The intervertebral space between T3 and T4 was identified, and a 32-gauge needle attached to a 10 μL Hamilton syringe was used to inject 0.5 μL of 1% w/v LPC (Sigma-Aldrich, L1381) into the ventral column at a rate of 0.25 μLmin-1 for 2 min. The needle was left for 2 min following the injection to avoid back flow. After suturing, mice were placed in a thermally controlled environment for recovery.

### Spinal cord tissue isolation

Anaesthetized mice were transcardially perfused with 15 mL of phosphate-buffered saline (PBS). Dissected spinal cords were fixed in 4% paraformaldehyde (PFA) at 4°C overnight. Tissue was cryoprotected in 30% w/v sucrose solution for 72h then frozen in FSC 22 frozen section media (Leica). Coronal sections were cut with a cryostat and stored at -20°C until analysis.

### Immunofluorescence staining

Slides were permeabilized with 0.2% TritonX-100 and blocked with a 10% horse serum-containing solution. Primary antibody incubation was conducted overnight in phosphate-buffered saline (PBS) containing 0.1% cold fish stain gelatin and 0.1% Triton X-100. Slides were then stained with TrueBlack Lipofuscin Autofluorescence Quencher (Biotium) for 2 min, and incubated with secondary antibody and 1 µgmL-1 of DAPI for 1h before mounting.

Antibodies used were anti-mouse myelin basic protein (MBP, BioLegend, PA1-10008), anti-mouse periostin (POSTN, R&D Systems AF2955), anti-mouse α smooth muscle actin (αSMA-Cy3, Millipore, C6198), anti-mouse fibronectin (FN1, Millipore, AB2033), anti-CD45 (Thermofisher, MA5-17687), rabbit anti-mouse pan-laminin (gift from L. Sorokin, University of Münster), anti-mouse/human PDGFRβ (R&D Systems, AF1042), anti-mouse Iba1 (Wako, 019-19741), anti-mouse CD45-AF488 (Biolegend, 103122), rabbit anti-collagen 1a1 (Col1a1, Invitrogen, PA1-26204), anti-mouse GFAP (Biolegend, PCK-591P), anti-mouse NFH (RPCA-NF-H), anti-mouse PDGFRα (R&D Systems, AF1062), anti-mouse Olig2 (Millipore, ab9610), anti-green fluorescence protein (GFP, Aveslab, GFP-1020), anti-mouse sulfatide O4 (R&D Systems, MAB1326), anti-mouse arginase-1 (Arg1, BioLegend, 678802), and anti-mouse MHCII (Thermofisher, 13-5321-82). Secondary antibodies from Jackson ImmunoResearch were AF488 donkey anti-mouse IgM, AF488 donkey anti-rabbit IgG, AF488 donkey anti-rat IgG, AF488 donkey anti-goat IgG, cyanine Cy3 donkey anti-goat IgG, cyanine Cy3 donkey anti-chicken IgY, AF647 donkey anti-rat IgG, and AF647 donkey anti-rabbit IgG.

### Confocal immunofluorescence microscopy and analysis

Images were collected with a Leica TCS Sp8 laser confocal microscope. Equal laser, gain and offset settings to maximize contrast and minimize saturation were consistently used for all samples within each set of experiments. Leica Application Suite X, ImageJ, and QuPath 0.3.1 were used for image acquisition and analysis. Maximum-intensity projections were created for each channel/marker. All images were blinded prior to analysis. ROI for lesions were drawn using DAPI or MBP. Volume analysis of ventral white matter lesions was done using MBP immunostaining of serial sections to manually trace the region of demyelination largely devoid of myelin, or containing intense fragmented MBP signal, per section, across the coronal sections from each mouse. Fibroblast regions were defined using threshold of PDGFRβ-Tdtomato or PDGFRβ immunostaining. Proportion of lesion occupied by fibroblasts was calculated by dividing the PDGFRβ-Tdtomato or PDGFRβ immunostained area by the MBP derived lesion area for the respective sections. Threshold intensity was determined using at least 3 images. The Analyze Particles function was then used to create a mask and to quantify the positive signals in each ROI using size exclusion of 2-infinity. Threshold intensity, size exclusion, and circularity settings for particle analysis were kept constant across all samples for each experimental set. Oligodendrocytes were counted manually using Olig2 and DAPI as a guide. When analyzing lesion spread, samples were excluded if more than 20% of sections were unable to be analysed due to tearing, folding, or other loss.

### Cell cultures and analyses

For fibroblasts, meninges from P0-2 CD1 pups were collected in RPMI and digested for 18 minutes at 37°C in 1 μgmL-1 DNase I (Sigma) and 3.7 μgmL-1 collagenase D (Sigma) triturating every 6 minutes. Meninges were centrifuged at 300xg for 3 minutes and cells were resuspended in growth medium (DMEM (Gibco) supplemented with 10% FBS, 1% non-essential amino-acids (NEAA), 1% GlutaMAX, 1% sodium pyruvate, and 1% penicillin-streptomycin (all from Gibco). Cells were incubated in T-75 flasks at 37°C with 5% CO2. Medium was replaced every 3 days and cells were passaged when 90% confluent.

Mouse OPCs were cultured from cortices of P0 CD1 pups as previously described^21^. Mouse bone marrow-derived macrophages (BMDMs) were generated as detailed elsewhere^32^.

For co-cultures, OPCs were added at a density of 31,250 cells/cm^2^ to poly-L-lysine coated 96-well plates. Meningeal fibroblasts were added at particular densities as listed in Results. OPCs and meningeal fibroblasts were added together, or one added 2h before the other as defined in the experiment, in 100 µL of OPC differentiating medium. Cocultures were incubated at 37°C with 8.5% CO_2_ for 72 hours. Cells were fixed and anti-O4 primary antibody was added in Licor antibody dilution buffer (1:250) overnight at 4°C. Donkey anti-mouse IgM AF488 secondary antibody was then added (1:200) followed by permeabilization with 0.02% Triton-X100 in PBS. Cells were further incubasted overnight at 4°C with anti-PDGFRβ (1:50; R&D systems;AF1042) and anti-MBP (1:100; Abcam; ab7349). When indicated, fibroblasts were stained with CellTracker Deep Red (Thermofisher; C34565) before plating. After secondary antibodies (1:200) exposure, cells were suspended in PBS with 1 μg/mL DAPI until imaging.

For analyses of stained plates, these were imaged using a ×10/0.5 NA air objective on an ImageXpress Micro XLS High-Content Analysis system (Molecular Devices) as previously described^24,30^. Nine–12 fields of views (FOVs) were imaged per well. Multiwavelength cell scoring analysis in the MetaXpress High-Content Image Acquisition and Analysis software (Molecular Devices) was used to quantify cell survival from the fluorescence microscopy images gathered. For fold change values, all numbers were divided by the mean of the control samples. For the representative images shown, each channel/marker for the sample was merged and displayed using pseudocolours.

### Transwell migration assays

BMDMs were plated in 24 well plates at a density of 260,000 cells/cm^2^ overnight at 37°C with 8.5% CO_2_. Medium with 1% FBS was added for 24h prior to experiments. Meningeal fibroblasts (150,000 cells/cm^2^) were added to transwell inserts (Corning, 29442-120) for 16h, then fixed and stained. The upper chamber was cleaned using a cotton swab, inserts were transferred to hematoxylin for 15 minutes, washed, mounted and imaged on an Olympus VS110 slide scanner on a 10x/0.4 NA objective and VS-ASW-S5 (V2.9) with Batch Converter software.

### CosMx spatial molecular imaging and data analysis

FFPE sections as prepared above were sent to NanoString for CosMx platform analysis. Sample processing, staining, imaging and cell segmentation were performed as previously described^33^. The 1000-plex CosMx Mouse Neuro probe panel was used along with markers for morphology and cell segmentation. The CosMx optical system has an epifluorescent configuration based on a customized water objective (13×, NA 0.82), and uses widefield illumination, with a mix of lasers and light-emitting diodes to image DAPI, Alexa Fluor-488, Atto-532, Dyomics Dy-605 and Alexa Fluor-647, as well as removal of photocleavable dye components. Nine 0.8 um Z-stack images of each FOV were acquired and then fluorophores on the reporter probes were UV cleaved and washed off with Strip Wash buffer. This procedure was repeated for the remaining 15 reporter pools, and the 16 cycles of reporter hybridization-imaging was repeated 8 times to increase RNA detection sensitivity. Raw image processing and feature extraction were performed using an in-house SMI data processing pipeline which includes registration, feature detection, and localization^33^. A machine learning algorithm cell segmentation pipeline was used to assign transcripts to cell locations and subcellular compartments as previously described^34^.

Data from the CosMx platform was formulated into an object for Seurat v5^35^ in R. Data from 2 LPC at 14 days post-lesion (dpl) and 3 Sham brain sections with FOVs from the right hemisphere were used. Data was normalized using SCTransform v2. Unsupervised analysis and generation of a final UMAP with 20 PCs and resolution = 0.7 was done. Harmony^36^ was used to integrate ‘Group’ data (LPS or Sham). Robust Cell Type Decomposition (RCTD) was used to map the cluster profiles to the Azimuth mouse motor cortex reference map (azimuth.hubmapconsortium.org). The following cell types were identified: astrocytes, endothelial cells, neurons, macrophages, oligodendrocytes, oligodendrocyte precursor cells (OPCs), vascular leptomeningeal cells (VLMCs) and pericytes. A separate fibroblast identity was created expressing *Col1a2* and *Pdgfrb*, and for OPCs expressing *Pdgfra* and *Sox6*. FindAllMarkers with default Wilcoxon rank sum test was used to identify distinguishing markers for each cluster. DoHeatmap with downsampling to 1000 was used to generate a heatmap with top 5 genes for each cluster. Data from macrophages, fibroblasts and OPCs from LPC sections were subset from the main dataset and used with the package CellChat v2 to assess ligand-receptor interactions^37^. To assess significant ligand-receptor interactions the computeCommunProb command was used with the following parameters (type=”truncatedMean”, distance.use=TRUE, contact.dependent =TRUE, contact.range=50).

### Analysis of mouse single nuclei dataset from LPC lesions

Fibroblast expansion and activation were validated in a publicly available LPC-lesioned tissue dataset (Melchor et al. accession number GSE278643). Briefly, processed data were directly downloaded, and all quality control, dimensionality reduction, and clustering were performed using Seurat (v5.1.0) in R (v4.4.1), unless otherwise noted. All samples (naïve spinal cord tissue (n=2), LPC and internal non-lesion controls 5dpl (n=3, n=1 respectively), 10dpl (n=2, n=1 respectively), and 20dpl (n=3, n=1 respectively) were utilized. Quality control and filtering were consistently applied to all samples and took a dual universal and cluster-by-cluster approach. Briefly, universal standards for retaining quality nuclei were as follows: nUMI>500, nGene>300, mitochondrial gene percentage<5%, and ribosomal gene percentage<5%. We clustered each sample (50 PCAs, 0.4 resolution) and applied cluster-by-cluster standards for retaining quality nuclei samples such that nuclei with nGene<3MADs were retained. Doublets were calculated using scDblFinder(v1.8.0) with standard settings and excluded during filtration. Post-filtration, data were merged using the merge function from Seurat. The merged dataset was log normalized and variance stabilized using SCTransform (v2) ^38,39^. Principal component analysis (PCA) was performed using the top 2,000 most variable genes calculating 75 PCA dimensions. Clustering was then performed via FindNeighbors and FindClusters with the 75 PCA dimensions and a resolution of 0.8. The count matrix was log normalized using the NormalizeData function. All cell populations were annotated based off well-known and calculated marker genes using the FindAllMarkers function. Vascular and mesenchymal cells (*Pdgfrb, Cfap43, Pecam1, Cspg4, Col1a2*) were subset based off these marker genes. The subset vascular/mesenchymal dataset was reclustered (40 PCs and resolution = 1.5) and filtered to remove low quality nuclei, defined as expressing multiple cell markers, not captured by initial filtration (number of filtered nuclei = 87). The count matrix was then renormalized using the NormalizeData function, and clusters were manually annotated based on the expression of cell type-specific genes (Supplementary Fig 1). Fibroblasts (*Col3a1, Col1a2, Col1a1, Bnc2, Dcn*, and *Foxp2)* were then subset, reclustered (20 PCs and resolution = 1.0), and reannotated based off previously established marker genes^9,40^ to enable in-depth characterization of the fibroblast populations. Data visuals were generated using the scCustomize(v3.2.2) and ggplot2(v4.0.1) packages, unless otherwise noted.

### snRNAseq abundance analysis

To rigorously assess biological shifts, we calculated the statistical significance of differential abundance for every annotated cell cluster in injury conditions (Nonlesion, 5 dpl, 10 dpl, and 20 dpl) versus baseline naïve controls using the propeller function from the speckle package (v1.10.0)^41^. Propeller accounts for the biological variability between independent biological samples (n=2-3 for the dataset). Briefly, snRNAseq metadata comprised of cell type annotations, sample IDs, and experimental conditions were extracted from the Seurat object. Relative proportions of each cell type were calculated for each sample. The proportions were then transformed using an arcsine square root transformation and a linear model was fitted to these transformed values per pairwise test against naïve controls (T-tests). P-values were adjusted using the Benjamini-Hochberg false discovery rate (FDR). Annotated cell clusters with an adjusted p-value < 0.1 were trending significance, and cell clusters with an adjusted p-value < 0.05 were considered significant. A delta value was calculated to indicate the biological difference in relative proportions for each test and was utilized for plots to highlight the largest biological shifts. Full results are reported in Supplementary Table 1.

### CellChat cellular signaling analysis of LPC snRNAseq data

Fibroblasts, microglia/macrophages and OPCs at 5 and 10 dpl were subset from the LPC snRNAseq dataset and analyzed with CellChat (v2) to assess communication potential between the three cell types, post-demyelination. To assess significant ligand-receptor interactions the computeCommunProb command of CellChat was used with the following parameters: type=”truncatedMean”, trim = 0.05, population.size=TRUE. Native CellChat and custom ggplot2(v4.0.1) plots utilizing the same data were used to visualize the analyses. For several data visuals, only top signaling family pathways or Ligand-Receptor pairs are shown, however all data are included in Supplementary Table 2. The selection of top pathways is detailed in figure legends but was defined by highest communication probability (weight/strength).

### Analysis of human single nucleus RNA dataset from MS patients

Fastq files from Absinta et al. (accession number GSE180759) were downloaded and demultiplexed using 10x Genomics Cellranger version 5.0 pipeline. The final expression matrix with gene counts from n=3 controls and n=5 patients was analysed using the bioinformatics package Seurat v5^35^. Metadata defining tissue area was included in the object. Before running QC metrics the expression matrix comprised 34,131 features and 134,932 cells. Data was filtered for parameters: gene present in > 3 cells and cells with < 150,000 nCount_RNA and percent of mitochondrial genes < 10%. Post filtering, the expression matrix contained 33,454 features and 128,343 cells. Data was normalized and scaled using the SCTransform v2 function with percent.mt being regressed out. A PCA-reduction was performed and 30 significant dimensions were considered to generate a UMAP with all cells in the dataset. Clusters were determined using the FindClusters function which implements the Louvain algorithm for modularity optimization and with resolution of 0.3. Cluster annotation was done manually based on the expression of lineage specific hallmark genes. Differentially expressed genes for one cluster (versus all cells in other clusters) was determined by the default Wilcoxon rank sum test. *PDGFRB*+ cells from the initial fibroblast cluster were then subset and re-clustered. Data from microglia/macrophages, OPCs and *PDGFRB*+ fibroblasts were subset from the main data and used with the package CellChat2 v2 to assess ligand-receptor interactions^37^. To assess significant ligand-receptor interactions the computeCommunProb command was used with the following parameters (type=”truncatedMean”, trim = 0.05, population.size=TRUE).

### Statistical analysis

Data was collated using Microsoft Excel. Graphs were generated using GraphPad Prism 8. Data shown are the individual data points, each point on a bar graph represents a biological (in vivo) or technical replicate (in vitro), and the mean. Sample sizes were similar to those previously reported^30,42,43^. Experimental groups were randomly assigned. Data collection and analysis were not performed blind to the conditions of the experiment (unless otherwise stated), as all image and data analyses were completed with the same acquisition conditions and analysis thresholds. Statistical tests are listed in figure legends.

## Results

### Fibroblasts are localized within demyelinated lesions

Although restricted to CNS borders in homeostasis, fibroblasts are found in the parenchyma after neural injury^3,5–11^. However, their functional connectivity and contribution to remyelination remain poorly understood. The LPC model of demyelination allows precise spatial and temporal dissection of pathophysiology in lesions^43,44^. Injection of LPC into the ventral spinal cord white matter of mice results in a focal lesion characterized by loss or disrupted MBP stain, swollen axons based on neurofilament heavy chain staining (NFH), and GFAP-positive reactive astrocytes (Fig 1a). In PDGFRβ^TdTom^ transgenic mice commonly used to study fibroblast responses in peripheral organs^45–47^, we found that TdTomato appeared to be in juxtaposition with microglia/macrophage markers CD68 and Iba1 within LPC lesions (Fig 1b). As anticipated, we observed in LPC lesions that PDGFRβ immunoreactivity using an anti-PDGFRβ antibody was associated with but did not largely overlap with the TdTomato signal (Fig 1c) as the latter in PDGFRβ^CreERT2^:Rosa26^TdTom^ mice was not directly conjugated to PDGFRβ itself. We found that PDGFRβ immunoreactivity was further associated with collagen type 1 alpha 1 (Col1a1), a major ECM component deposited by pathological fibroblasts, and the myofibroblast marker smooth muscle actin (SMA) (Fig 1d).

**Fig. 1.**
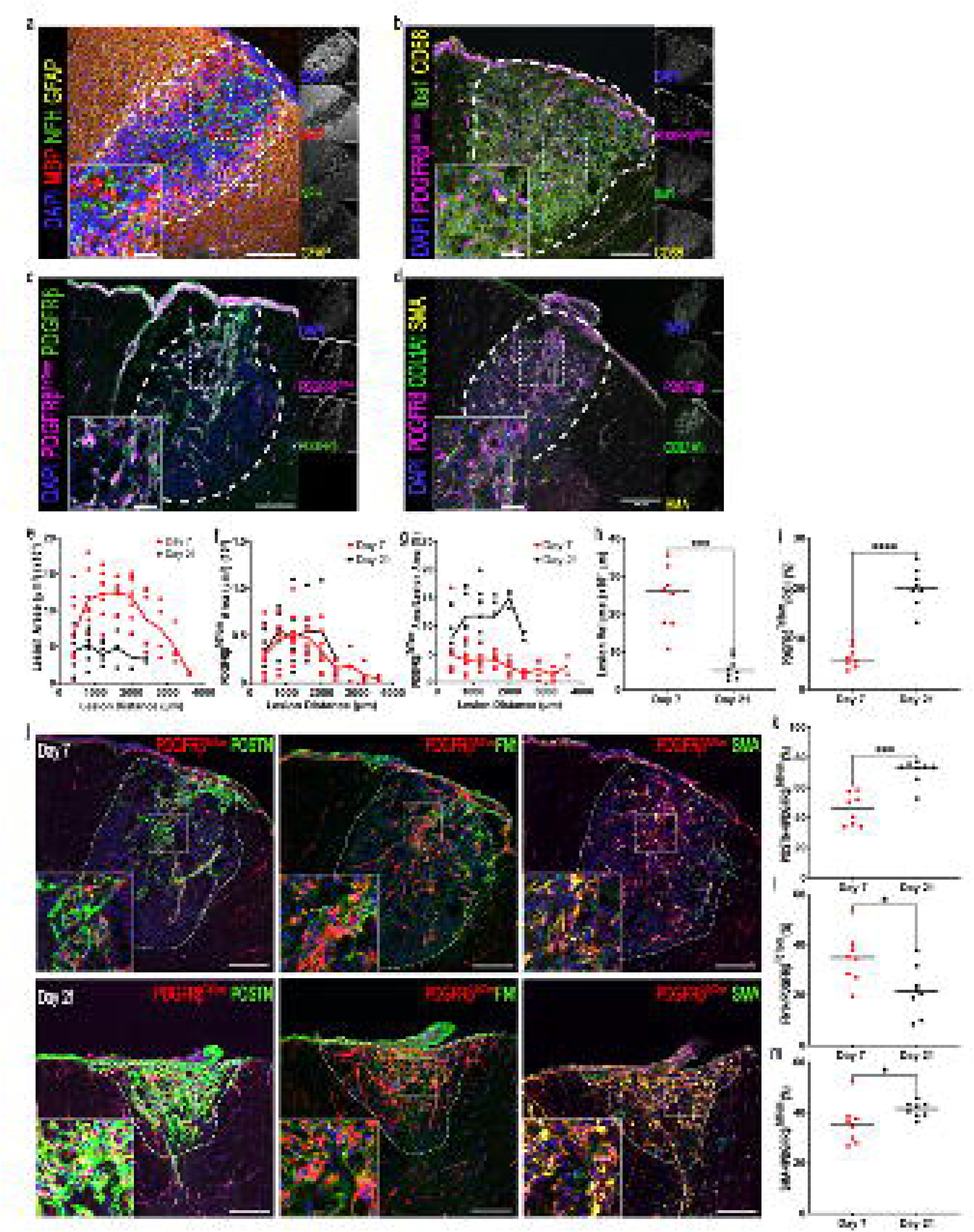
Fibroblasts localize in LPC demyelinated spinal cord white matter. (**a-d**) Immunofluorescence images of day 7 LPC lesions of 6-10 weeks old mice labeled with: (**a**) DAPI, MBP, NFH and GFAP in wildtype mice; (**b**) DAPI, CD68, Iba1 and TdTomato (TdTom) in PDGFRβ^TdTom^ mice; (**c**) DAPI, PDGFRβ antibody labeling and PDGFRβ in PDGFRβ^TdTom^ mice; (**d**) DAPI, PDGFRβ, COL1A1 and SMA in wildtype mice. Scale bar = 100 μm, (**e-i**) Comparison of day 7 and day 21 lesion, where each dot represents data per section per animal from the beginning of the lesion (zero distance) for (**e**) lesion area, (**f**) TdTom+ area, or (**g**) TdTom+ expressed as % of lesion area. Solid line in e-g represents the mean. The area under the curve (AUC) across the sections (thus volume) for individual mice are shown for (**h**) lesion size, and (**i**) TdTom region of interest (ROI). (**j**) Representative images of day 7 and 21 LPC lesions labeling for PDGFRβ^TdTom^ and (left) periostin (POSTN), (middle) fibronectin (FN1), and (right) SMA. Scale bar = 100 μm. (**k-n**) Graphs comparing the proportion of day 7 and 21 lesions positive for (**k**) POSTN, (**l**) FN1, (**m**) SMA. Data was acquired from two experiments; *n*=8 per experimental group. Significance is indicated as \**P*<0.05, \*\**P*<0.01, \*\*\**P*<0.001, \*\*\*\**P*<0.0001; unpaired *t*-test, comparing day 7 and day 21 LPC lesions. Data presented as the mean.

Fibroblast responses were studied 7 and 21 days post-lesion (dpl), periods when tissue regenerative responses are robust and when they are concluding, respectively^22,43^ (Fig 1e-i). While the total lesion volume was reduced from day 7 to 21 (Fig 1e, h), the total TdTomato-positive volume was unchanged between 7 to 21 dpl (Fig 1f) resulting in a greater proportion of the lesion occupied by fibroblasts at 21 dpl (Fig 1g, i).

To investigate whether fibroblast activation changed over time, we evaluated expression of SMA, as well as ECM components fibronectin (FN1) and periostin (POSTN) that are associated with a myofibroblast phenotype^9,48^ (Fig 1j). Roughly 40% of TdTomato-positive fibroblasts overlapped with each marker at 7 dpl with SMA and periostin increasing at 21 dpl and fibronectin decreasing at 21 dpl (Fig 1k-m).

### Single-nucleus RNA sequencing defines fibroblasts in LPC lesions

We utilized an in-house single nucleus RNA sequencing (snRNAseq) (GSE278643) dataset of LPC lesions at 5, 10 and 20 dpl (Fig 2; Supplementary Fig 1, 2). From all identified cell types (Supplementary Fig 1a), vascular and mesenchymal cells in LPC lesions were clustered and annotated into five groups: fibroblasts (*Col3a1, Col1a2, Col1a1, Bnc2, Dcn*, *Pdgfra* and *Foxp2*)^9,40,49^, endothelial cells (*Flt1, Pecam1, Ly6c1* and *Adgrl4*)^49^, pericytes (*Atp13a5, Abcc9, Cspg4* and *Kcnj8*)^49,50^, smooth muscle cell (SMC) (*Acta2, Myh11, Lmod1, Myom1* and *Tagln*)^49^, and ependymal cells (*Foxj1, Cfap54, Dnah12* and *Cfap299*)^51,52^ (Supplementary Fig 1b-d). *Pdgfrb* was variably expressed in fibroblasts, pericytes and SMCs as has been previously reported in transcriptomic studies^49^. Fibroblasts were expanded in LPC lesions at 5 and 10 dpl (Supplementary Fig 1e-g; Supplementary Table 1). Fibroblasts constituted ∼75% of the vascular and mesenchymal cells at 5 and 10 dpl, and approximately 60% at 20 dpl (Supplementary Fig 1b). Further evaluation of the non-fibroblast vascular and mesenchymal cell populations can be found elsewhere^53^.

**Figure 2.**
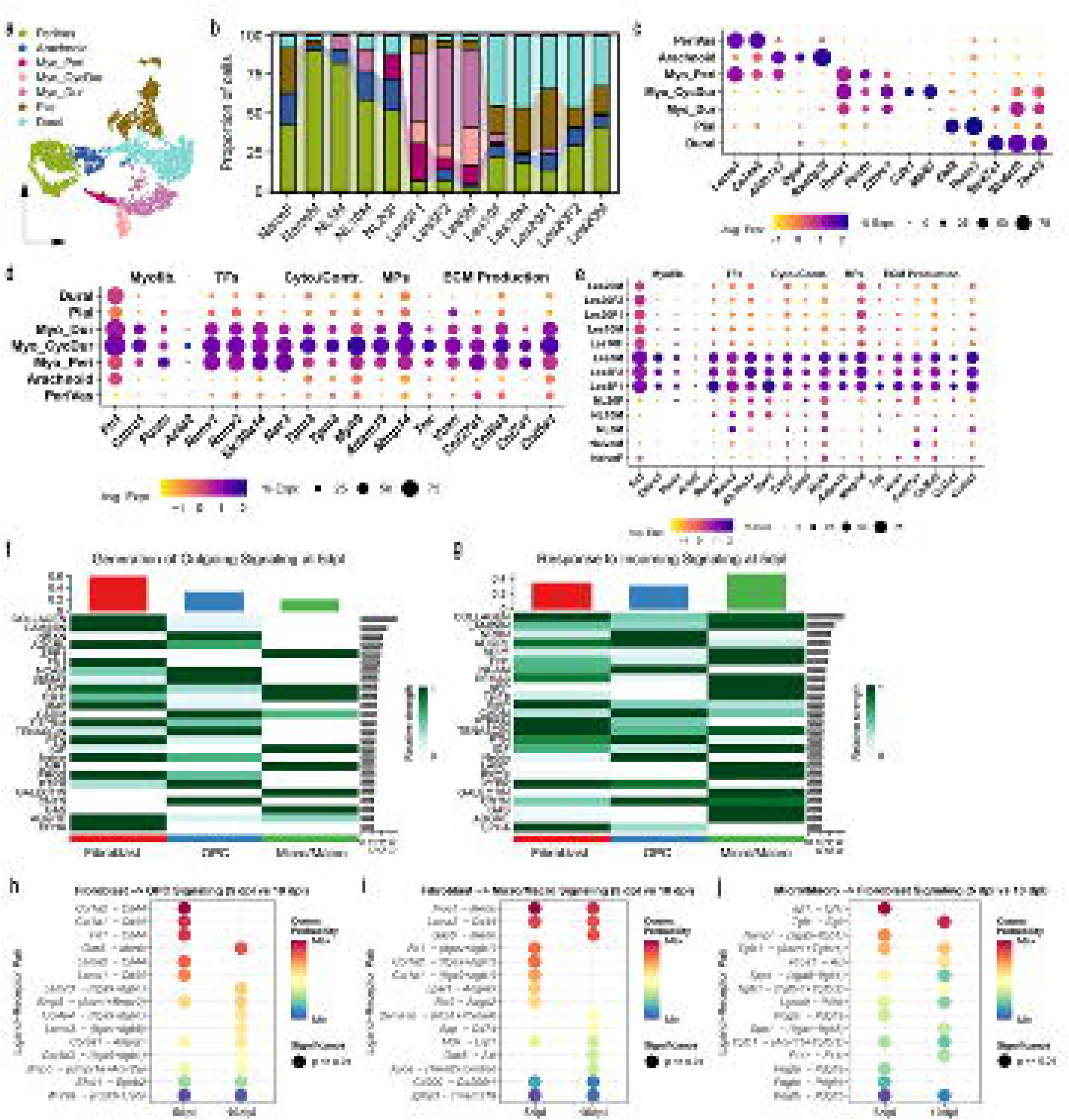
Single-nucleus RNA sequencing confirms the prominence of fibroblasts in LPC lesions. (**a**) UMAP plot of 3,443 annotated fibroblast subpopulations. (**b**) Bar plot showing the relative frequencies of fibroblast subpopulations in each experimental sample, highlighting the prominence of myofibroblasts at 5dpl. Abbreviations: F – female; M – male; NL – non-lesion, depicting a white matter spinal cord area several mm rostral from lesion, with no apparent injury; Les – LPC lesion. The numerals 5, 10 or 20 depict the days post-LPC when mice were killed for analyses. **(c)** Dotplot of annotated fibroblast subpopulations highlighting marker genes used for annotation. **(d)** Dotplot showing the expression of multiple activation and myofibroblast related genes across all fibroblast subpopulations. Myofib - myofibroblast-related; TFs - transcription factors; cyto/contr = cytoskeletal/contractile; MPs – metalloproteinases; ECM production - extracellular matrix production. **(e)** Dotplot showing the expression of activation and myofibroblast related genes in **(d)** across samples and timepoints. **(f, g)** Heatmaps showing the relative contribution of the generated outgoing **(f)** and incoming **(g)** signaling at 5dpl. **(h-j)** Bubble plots of top ligand-receptor (LR) pairs from fibroblasts to OPCs **(h)**, fibroblasts to microglia/macrophages **(i)**, and microglia/macrophages to fibroblasts **(j)** highlighting LR pairs associated uniquely with 5dpl, uniquely with 10dpl, and conserved LR pairs shared between the two timepoints. LR pairs were sorted by communication probability strength differential between the two timepoints. All significant interactions for 5 and 10 dpl are reported in Supplementary Table 2.

We subclustered fibroblasts and annotated them into 7 groups based off of previous literature^9,40^: perivascular (PeriVas: *Lama1*, *Col4a6*), arachnoid/arachnoid barrier (Arachnoid: *Aldh1a2*, *Dpp4*), pial-like (Pial: *Ebf2*, *Tenm2*) and dural (Dural: *Slc4a10*, *Slc47a1*) fibroblasts; and three myofibroblast populations including perivascular (Myo_Peri, *Lama1*, *Col4a6*), dural (Myo_Dur, *Slc4a10*, *Slc47a1*) and cycling dural (Myo_CycDur, *Mik67*, *Slc47a1*) myofibroblasts, that shared expression of myofibroblast markers *Cthrc1* and *Postn* (Fig 2a-c). The observed myofibroblast populations exhibited increased expression of myofibroblast-related transcription factors *Runx1* and *Runx2*, cytoskeletal and contractile *Nav3* and *Tpm3*, metalloproteinases *Adam19* and *Mmp14*, ECM proteins *Tnc* and *Vcan*, and several collagen genes including *Col27a1*, *Col6a3* and *Col7a1* (Fig 2d). All myofibroblast clusters were significantly expanded (Myo_Peri: OddsRatio: 21.8, Adj p-value < 0.001, Myo_Dur: OddsRatio: 50.4, Adj p-value < 0.001, Myo_CycDur: OddsRatio: 70.8, Adj p-value < 0.001) in LPC lesions predominantly at 5 dpl (Fig 2b; Supplementary Fig 1h-j; Supplementary Table 1). Beyond myofibroblasts found at 5 dpl, fibroblasts showed maintained expression of several myofibroblast related genes through 20 dpl (Fig 2e), indicating continued stress and corroborating reports in other conditions^53^.

Collectively, snRNAseq data show that fibroblasts expand rapidly, taking on a myofibroblast phenotype early after injury but lowering this phenotype as they persist into late stages of LPC injury.

### CellChat analyses of snRNAseq results

To determine interactions between fibroblasts and OPCs and microglia/macrophages within LPC lesions, we conducted CellChat analyses of the LPC lesion snRNAseq dataset at 5 and 10 dpl (Fig 2f-j; Supplementary Fig 2; Supplementary Table 2). We found that at both 5 and 10 dpl, fibroblasts largely dominated outgoing cellular signaling (Supplementary Fig 2b, d), while further dominating the incoming cellular signaling domain at 10 dpl (Supplementary Fig 2c, e). Fibroblasts at 5 dpl mainly participated in reciprocal communication with OPCs measured both by total counts and weight; however, fibroblasts also had strong outgoing communication to microglia/macrophages (Supplementary Fig 2b, d). The most notable deviation in communications between 5 and 10 dpl was the increase in ECM-receptor signaling from microglia/macrophages-to-fibroblasts and cell-cell contact from OPCs-to-fibroblasts (Supplementary Fig 2f, g).

Assessment of the top incoming and outgoing inferred fibroblast signaling pathways with OPCs and microglia/macrophages showed ECM-receptor pathway families such as COLLAGEN and LAMININ exhibited the strongest overall outgoing communication signaling at both 5 and 10 dpl (Fig 2f, g; Supplementary Fig 2h, i). Additionally, cell-cell contact pathways such as ADGRL and APP, and secreted signaling pathways such as BMP and PROS were also observed. Together, these results suggest a large communication potential between fibroblasts, microglia/macrophages, and OPCs. Importantly, cell-cell contact pathway families further suggest that fibroblasts, microglia/macrophages and OPCs are poised to communicate directly in the shared lesion space.

We investigated the ligand-receptor pairs underlying signaling from fibroblasts to OPCs at 5 and 10dpl (Supplementary Table 2). To highlight signaling changes over time we plotted top pairs (defined by probability strength) found exclusively at 5dpl, at 10dpl, and shared between the two timepoints (Fig 2h-j). Fibroblast derived ECM ligands (*Col1a1, Fn1, Lama2, Lamc1*) were commonly the strongest 5 dpl unique interactors predicted with OPCs (*Cd44)* and microglia/macrophages (*Itgb1, Itgav, Itga9*) (Fig 2h). Other fibroblast ligands were associated with regulation of OPC migration (*Col1a1, Ntn1, Tnc*), proliferation (*Ptn, Igf1, Fgf2*) and differentiation (*Lama2, Fn1, Hspg2*) (Fig 2h; Supplementary Table 2). As well, fibroblasts were a source of ligands important for immune function including recruitment (*Cxcl12, Cx3cl1*), adhesion (*Col1a1, Fn1, Icam1*) and immunomodulation (*Mdk, Tgfb2, Pros1*). Interestingly, immunomodulatory signals at 5 dpl are more associated with regenerative responses (*Ptn, Fgf1, Igf2*) while at 10 dpl they appear more pro-inflammatory (*Gas6, Spp1, App*) (Fig 2i). Signaling from microglia/macrophages towards fibroblasts at 5 dpl included signals relating to chemotaxis and polarization (*Pdgfa, Pdgfc, Igf1*) (Hung et al 2013), and at 10 dpl, signals that stimulate ECM production (*Tgfa, Tgfb1*) (Fig 2j; Supplementary Table 1). These results suggest that fibroblasts and microglia/macrophages reciprocally influence the other’s response in LPC lesions.

### Spatial transcriptomics of fibroblasts in LPC lesions

To verify spatial distribution of these fibroblast dynamics, we used the 1000-plex Mouse Neuroscience Panel with the in-situ single cell CosMx spatial platform from Nanostring. We analysed sham and LPC sections at 14 dpl, after peak fibroblast response and when remyelination is ongoing^43,54^ (Fig 3a; Supplementary Fig 3a, b). We identified 1828 cells from LPC and 1126 cells from sham controls in 12 clusters which were mapped to the Azimuth mouse motor cortex reference map (Fig 3b, c). This identified astrocytes (*Gja1, Slc4a4, Slc6a1),* oligodendrocytes (*Mag, Mog, Myrf*), OPCs (*Pdgfra, Cspg5, Vcan*), microglia/macrophages (*Csf1r, Hexb, Csf3r*), vascular leptomeningeal cells (VLMCs) (*Dcn, Vtn, Igfbp7*), endothelial cells (*Cldn5, Pecam1, Flt1*) and pericytes (*Rgs5, Vtn, Myl9*) (Fig 3d). A separate fibroblast identity was created for cells expressing *Col1a2* and *Pdgfrb* (Fig 3b-d). We identified a total of 84 *Col1a2* and *Pdgfrb* double-positive fibroblasts, 5 in sham controls and 79 in LPC lesions (Fig 3e, f). The top genes for fibroblasts aside from *Pdgfrb* and *Col1a2* were *Col3a1, Col1a1, Vtn* and *Dcn* (Fig 3d). Importantly, fibroblasts did not express the pericyte marker (*Rgs5*) or smooth muscle cell marker (*Tagln*) (Supplementary Fig 3c, d). Fibroblasts were 16 times more abundant in the parenchyma of LPC lesions at 14 dpl compared to sham (Fig 3e, f).

**Fig. 3.**
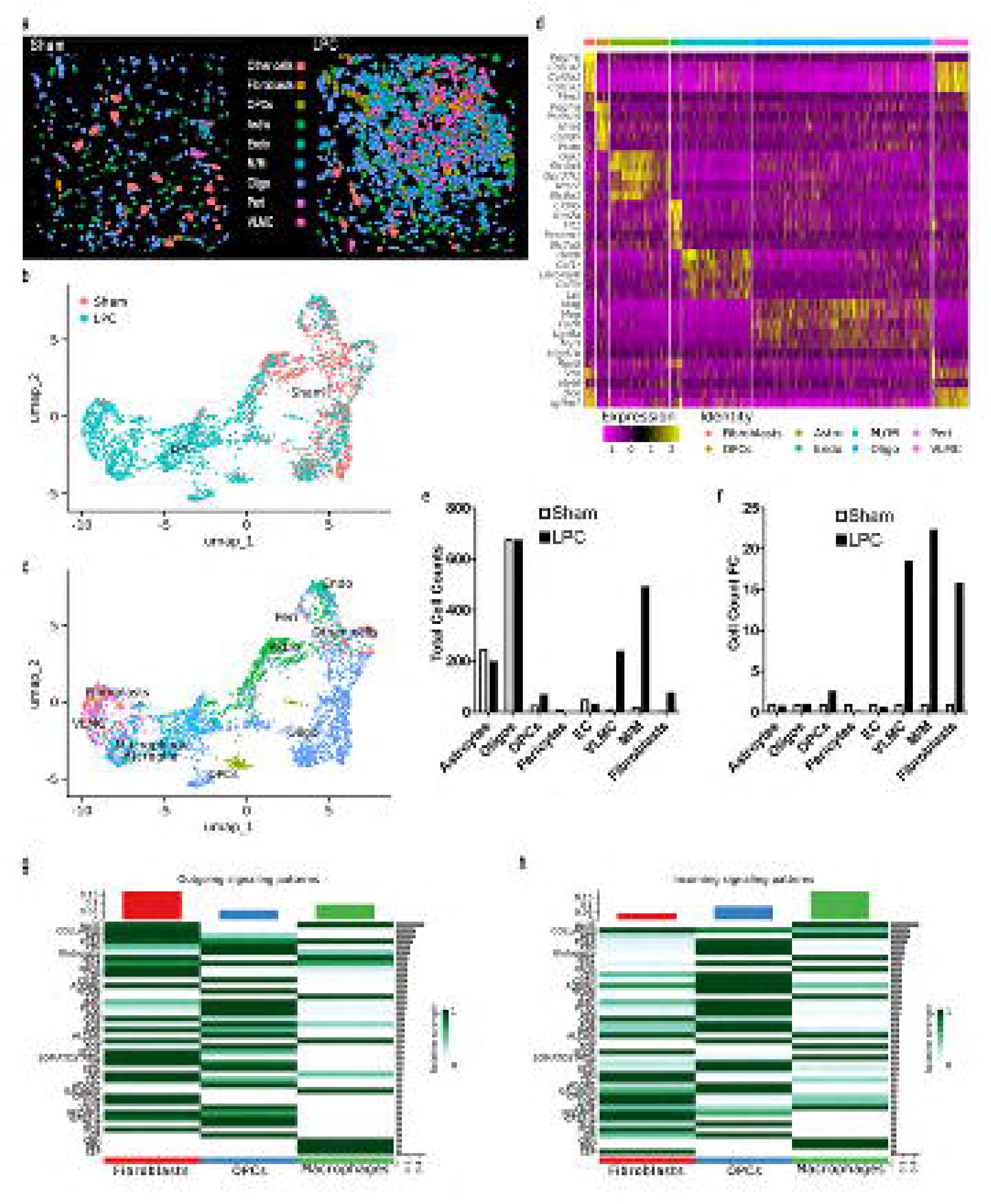
Spatial transcriptomics of fibroblasts in demyelinated lesions. (**a**) Representative images of annotated cells in sham and LPC demyelinated spinal cords at 14 dpl. Abbreviations: Astro – astrocytes; endo – endothelial cells; M/M – microglia/macrophages; oligo – oligodendrocytes; peri – pericytes; VLMC - vascular leptomeningeal cells. (**b, c**) UMAP visualization of 2954 cells from LPC and sham colored by (**b**) experimental group and (**c**) mapped cell clusters (20 principal components, resolution=0.7). (**d**) Heatmap of top 5 differentially expressed genes across annotated cell groups as determined using FindAllMarkers (default Wilcoxon Rank Sum test, min. logfc threshold = 0.25). (**e, f**) Bar graph showing (**e**) the total number of cells, and (**f**) the fold change in cell groups between LPC and sham groups. (**g, h**) Heatmap of (**g**) inferred outgoing and (**h**) incoming signaling pathways as assessed for the 3 cell groups. Color bar in the heatmap represents the relative signaling strength of a pathway across all cell groups (values are row-scaled). Top colored bar plot shows the total signaling strength of a cell group by summarizing all pathways displayed in the heatmap. The right grey bar plot shows the total signaling strength of a pathway by summarizing all cell groups displayed in the heatmap.

### Fibroblast interactions in LPC lesions

We utilized Cellchat^37^ ligand-receptor analysis to identify and compare general communication patterns in the spatial transcriptomic data between fibroblasts, microglia/macrophages and OPCs; we identified a robust fibroblast communication network in LPC lesions (Supplementary Fig 3e-i). Fibroblasts-OPC interactions were most common, followed by fibroblast-fibroblast interactions (Supplementary Fig 3e, g). However, the strongest interactions were between fibroblasts and microglia/macrophages (Supplementary Fig 3f, h). Fibroblast derived ligands affected receptors involved in signaling for phagocytosis (*Pros1-Axl, Pros1-Mertk*), macrophage polarization (*Spp1-Cd44, Apoe-Trem2/Tyrob*p), cell adhesion (*Col1a1-Itgav/Itgb8*) and OPC survival (*Ptn-Ptprz1, Bdnf-Ntrk2*) (Supplementary Fig 3j; Supplementary Table 3). The signals that fibroblasts were predicted to respond to influenced survival, proliferation, differentiation, and migration such as *Pdgfd-Pdgfrb, Fgf2-Fgfr2* and *Ptprs-Ntrk3* (Supplementary Fig 3k).

Categorization of ligand-receptor interactions into functionally related signaling pathways showed fibroblasts were the major driver of outgoing signaling in LPC lesions (Supplementary Fig 3l). Fibroblasts contributed to pathways such as apolipoprotein E (APOE), COLLAGEN, and FN1; microglia/macrophages to APOE, osteopontin (SPP1) and colony stimulating factor (CSF) pathways; and OPCs to pleiotrophin (PTN), Glutamate, and PDGF pathways (Fig 3g; Supplementary Fig 3l; Supplementary Table 3). Microglia/macrophages were the primary recipient of ligand-receptor signaling through pathways such as APOE and SPP1, and OPCs responded to PTN and FN1 (Fig 3h; Supplementary Fig 3m; Supplementary Table 3). Although fibroblasts were not identified as a dominant receiver of signaling through the cell types assessed, they were inferred to receive signaling predominantly from COLLAGEN and PDGF signaling pathways (Supplementary Fig 3i, m).

### Fibroblasts respond to microglia/macrophages

As fibroblasts are predicted to reciprocally communicate with microglia/macrophages in LPC lesions, we hypothesized that this might affect fibroblast expansion in the area of injury. Indeed, we found Iba1 and CD68 immunoreactivity representative of microglia/macrophages and PDGFRβ+ fibroblasts qualitatively associated with each other in day 7 LPC lesions (Fig 1b). Furthermore, Iba1+ microglia/macrophages began accumulating in the LPC lesion at day 3 before peaking at 7 days (Fig 4a-c) while PDGFRB+ fibroblasts also peaked at day 7, but there was no observable PDGFRβ+ immunofluorescence at day 3 (Fig 4a-d). Thereafter, there were no differences in the percent of the lesion positive for either PDGFRβ or Iba1 (Fig 4c, d). This temporal onset of microglia/macrophages and fibroblasts, and when considered together with CellChat analysis, suggests that microglia/macrophages might play a role in promoting fibroblast response in LPC. To evaluate this, we added meningeal fibroblasts to the upper compartment of a Boyden chamber with bone marrow-derived macrophages or medium alone in the lower chamber (Fig 4e). After 24 hours there was a 3-fold increase in the number of fibroblasts that migrated across the membrane when cultured with macrophages (Fig 4f, g). These results suggest a role for macrophages in the promotion of fibroblast elevation in CNS lesions.

**Fig. 4.**
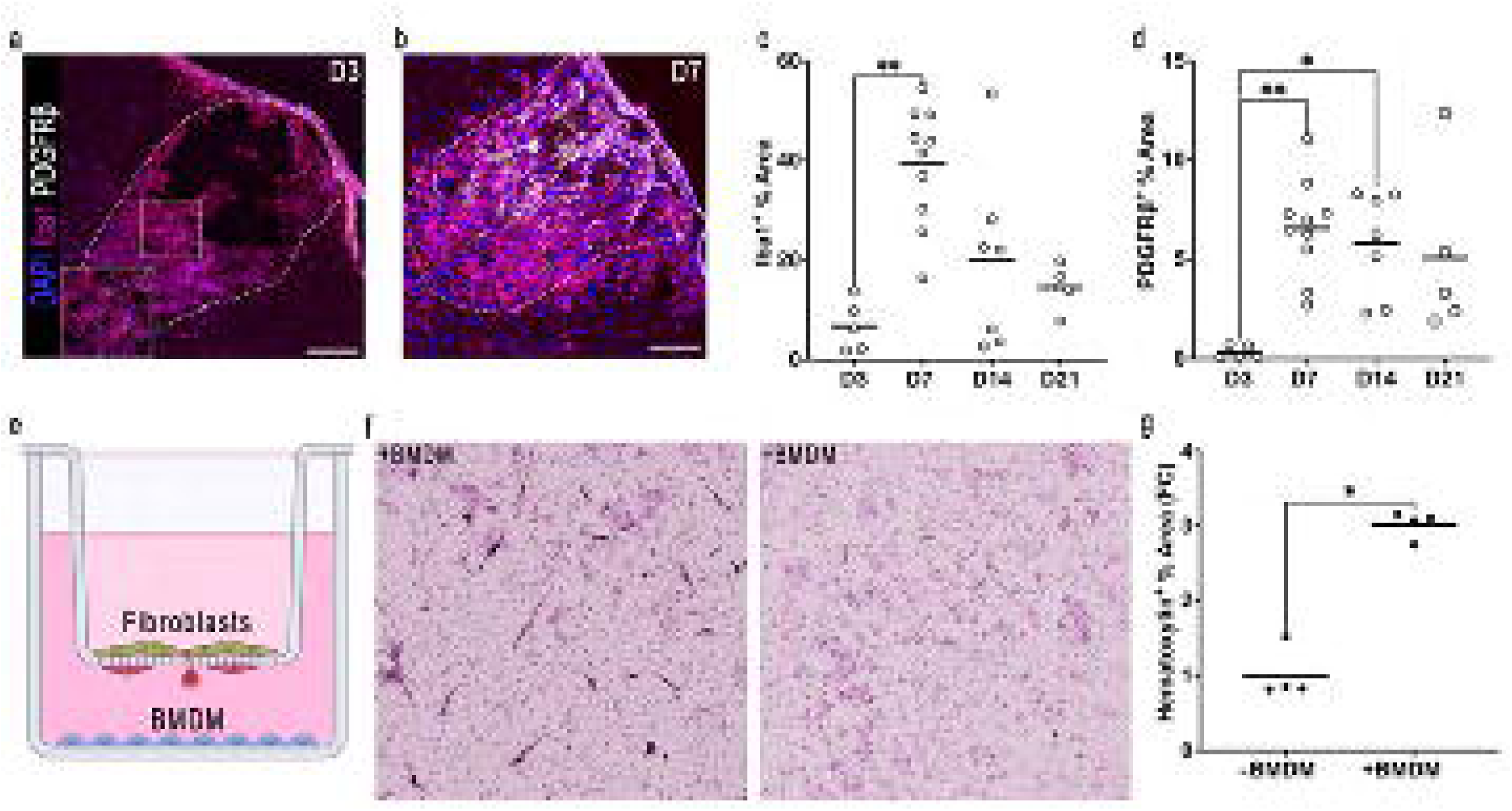
Post inflammation fibroblast response stimulated by microglia/macrophages. (**a, b**) Representative image of (**a**) day 3 and (**b**) day 7 LPC lesion labelling for DAPI, PDGFRβ, and Iba1. Scale bar = 100 μm. (**c, d**) Graphs indicating day 3, 7, 14, and 21 day for (**c**) Iba1 positive % area of lesion and (**d**) PDGFRβ positive % area of lesion. (**e**) Schematic of experimental design. Meningeal fibroblasts were added to upper compartment and BMDMs or medium alone added to the lower compartment. (**f**) Representative images of hematoxylin stained transwell inserts from BMDM and medium control groups. (**g**) Graph comparing the amount of hematoxylin stained meningeal fibroblasts migrated across the transwell membrane in BMDM and medium alone control. (**c, d**) *n*=5 (D3), 10 (D7), 7 (D14), 5 (D21); (**g**) *n* = 4 for each experimental group; Significance indicated as \**P*<0.05, \*\**P*<0.01, one-way ANOVA comparing time points (D3, 7, 14, or 21) (**c, d**); or two-tailed unpaired *t*-test (**g**). All data presented as the mean.

### Fibroblasts spatially exclude OPCs

In LPC lesions of wildtype mice, Olig2+PDGFRα+ OPCs were sparse in regions occupied by PDGFRβ+ fibroblasts (Fig 5a). This was corroborated in LPC lesions of NG2^CreER^:MAPT^mGFP^ mice in which a membrane-bound green fluorescent protein (GFP) is expressed under the tau (*Mapt*) promoter, the latter of which is enhanced in regenerating oligodendrocyte lineage cells ^20,21^ (Fig 5b, c). The reduction of OPCs in PDGFRβ+ areas was not due to reduced axonal preservation as we observed no difference in NFH+ axon density when comparing fibroblast occupied regions with the rest of the LPC lesion (Supplementary Fig. 4). These results suggest that fibroblasts restrict OPC localization in LPC lesions.

**Fig. 5.**
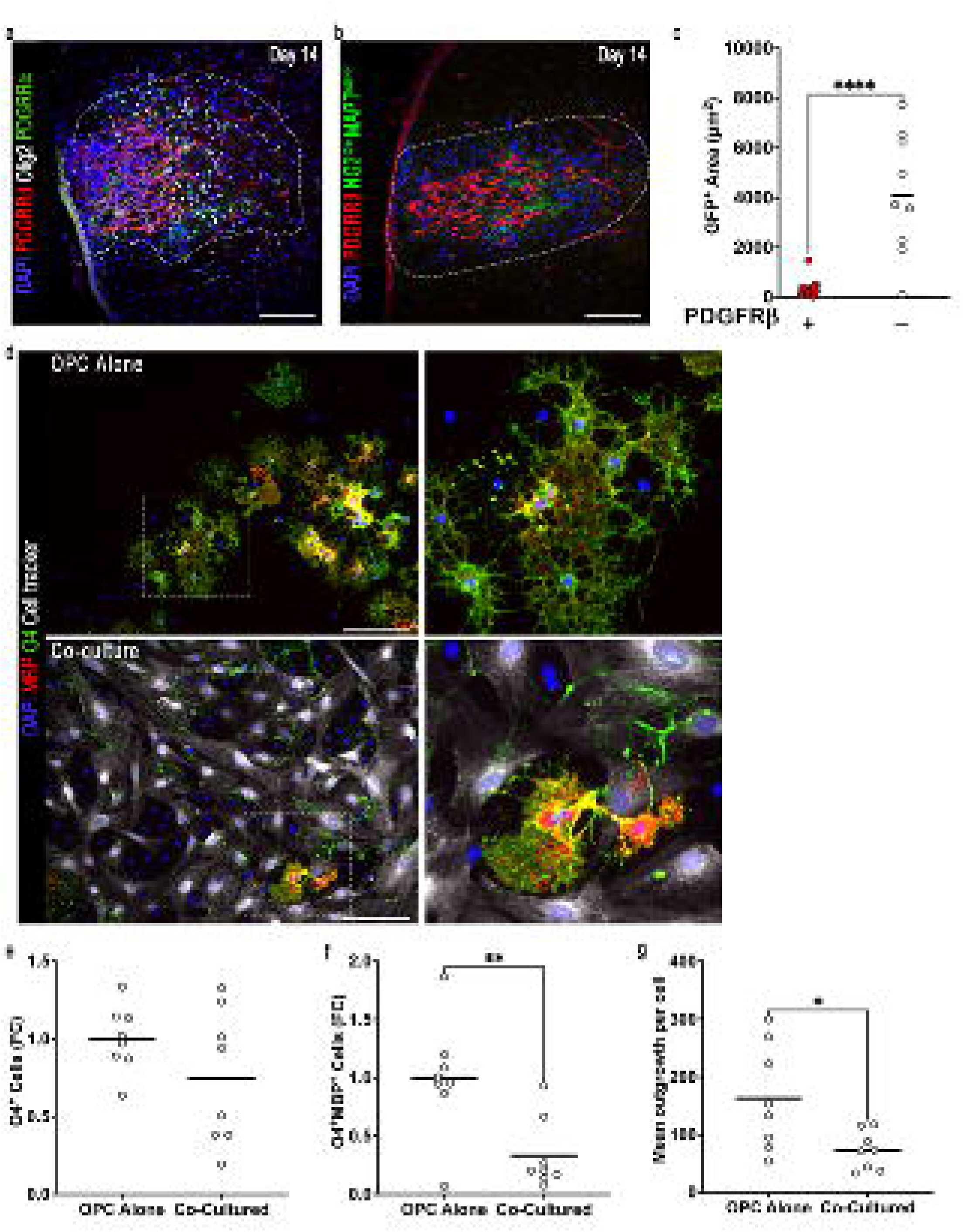
Fibroblasts spatially exclude OPCs in LPC lesions and in vitro. (**a**) Representative images of day 14 LPC lesion stained for PDGFRβ, Olig2, and PDGFRα. (**b**) Representative images of day 14 LPC lesion in NG2CreER:MAPTmGFP mice labelling for DAPI (blue), PDGFRβ using antibody (red), and NG2-GFP (green). (**c**) Graph comparing GFP (i.e. OPCs) positive area in day 14 LPC lesions in region of PDGFRβ+ (red) or PDGFRβ- cells. *n* = 9 each. Significance indicated \*\*\*\**P*<0.0001, two-tailed unpaired *t*-test. Data presented as mean. (**d**) Representative image of OPC alone (top, with magnified image of box at right) and OPC-fibroblast co-culture (bottom, with magnified image of box displayed at right) labelled for nuclei (blue), O4 (green), MBP (red), and CellTracker (fibroblasts, white). (**e-g**) Graphs indicating fold change in (**e**) O4+ OPCs, (**f**) O4+MBP+ mature oligodendrocytes, and (**g**) mean processes per cell comparing co-cultures with OPC alone controls. *n* = 2 independent experiments with 4 replicates for **e-g**. Significance indicated as \**P*<0.05, \*\**P*<0.01, two-tailed, unpaired *t*-test, Data presented as mean.

To test this further we seeded cultures of meningeal fibroblasts with OPCs (Fig 5d). No change in total O4+ OPC number was observed (Fig 5e) but a significant decrease in the number of mature O4+MBP+ oligodendrocytes in co-cultured wells was found (Fig 5f). Oligodendrocytes also had less complex morphology in co-cultures and they primarily occupied regions without fibroblasts (Fig 5d, g). Collectively, this data highlights the inhibitory effect of fibroblasts on OPC differentiation.

### Aging enhances fibroblast properties in LPC lesions

Age affects many biological processes including tissue regeneration^55^. Age is known to affect OPC and microglia/macrophage dynamics, and is a critical factor in remyelination potential^22,24,56,57^. To test how age affects the fibroblast response to CNS injury and the downstream effects, young (6-10 week) and middle-aged (48-52 week) mice were injected with LPC in the ventral column of the spinal cord. Tissue was collected 7, 14 and 21 dpl and immunostained for MBP and PDGFRβ (Fig 6a). LPC lesions in middle-aged mice were significantly larger at 14 and 21 dpl compared to the younger animals (Fig 6b, c). As well, a greater proportion of the LPC lesions in middle-aged mice were positive for PDGFRβ+ fibroblasts at 7 and 21 dpl when compared to lesions in young mice (Fig 6d).

**Fig. 6.**
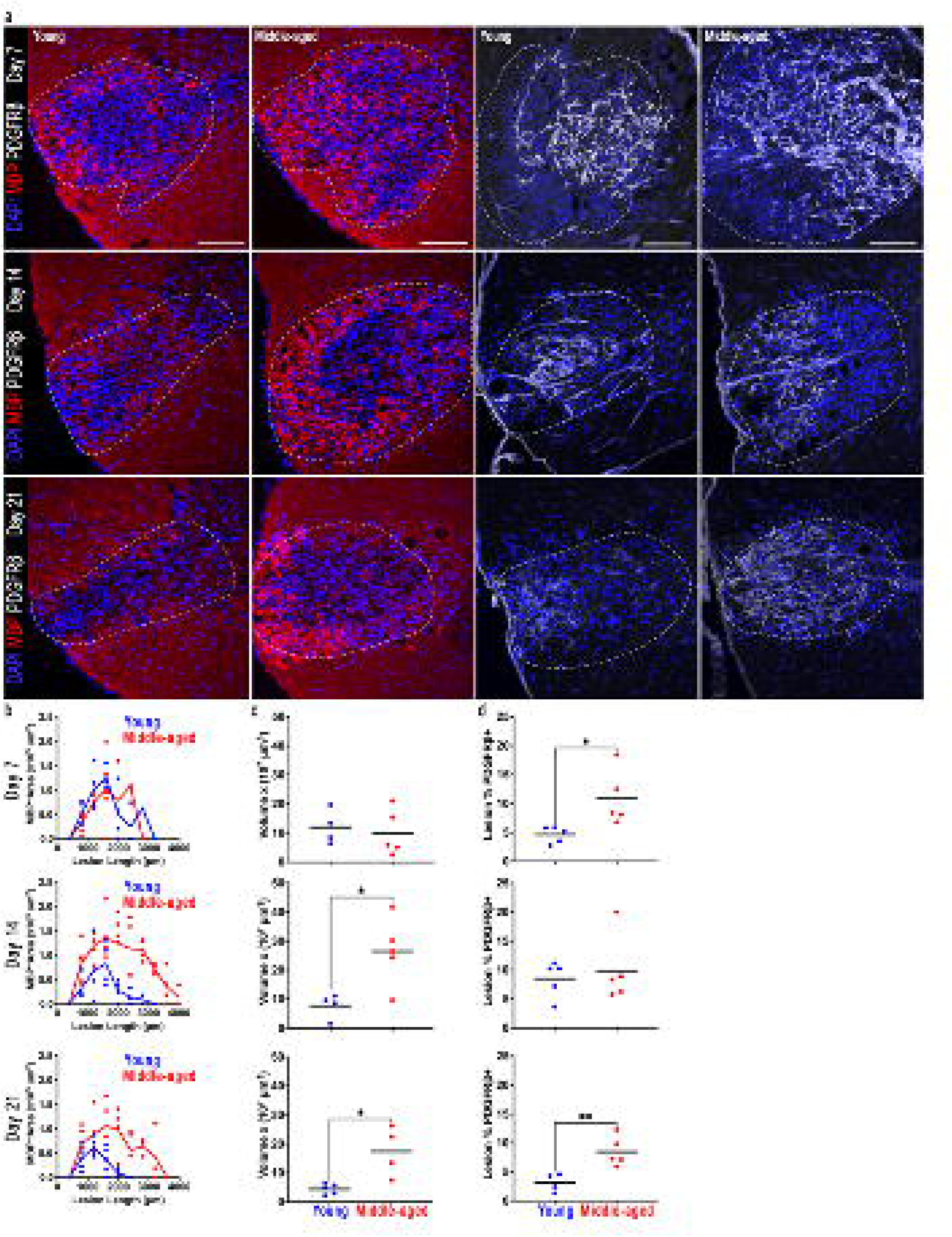
Age exacerbates fibroblast response to demyelination. (**a**) Representative immunofluorescence images of 7, 14 and 21 day LPC lesions in young (6-10 week old) and middle-aged (48-52 weeks) mice labeled with DAPI and MBP (red) or PDGFRβ (white). Lesion is indicated with white dashed line. (**b-d**) Graphs comparing the extent of demyelination (MBP-minus) and PDGFRβ+ fibroblasts in LPC-spinal cord lesions at days 7, 14 and 21 in young and middle-aged mice, where each dot in panel (**b)** represents data per section per animal from the beginning of the lesion (zero distance); the solid line represents the mean; the data across each animal is then used to generate the corresponding volume of lesion (**c**) and the % of lesion covered by PDGFRβ+ immunoreactivity (**d**). For young group, *n* = 4 (D7), 4 (D14), 5 (D21); for middle-aged group *n* = 5 for each time point. Significance is indicated as \**P*<0.05, **P<0.01; unpaired, *t*-test, comparing young and middle-aged groups. Data is presented as the mean.

We investigated the effect of age on the expression of phenotypic markers in fibroblasts in LPC lesions. Middle-aged LPC fibroblasts at day 7 displayed elevated levels of myofibroblast marker SMA compared to young (Fig 7a). While no differences in Col1a1 or laminin were seen at 7 dpl, these were elevated at 21 dpl (Fig 7b, c). Finally, we found fewer olig2+ oligodendrocyte lineage cells within the fibroblast-occupied regions of middle-aged compared to young mice (Fig 7d). Overall, elevated ECM levels in middle-aged compared to young mice corresponded with a greater reduction in oligodendrocyte lineage cells in lysolecithin injury.

**Fig. 7.**
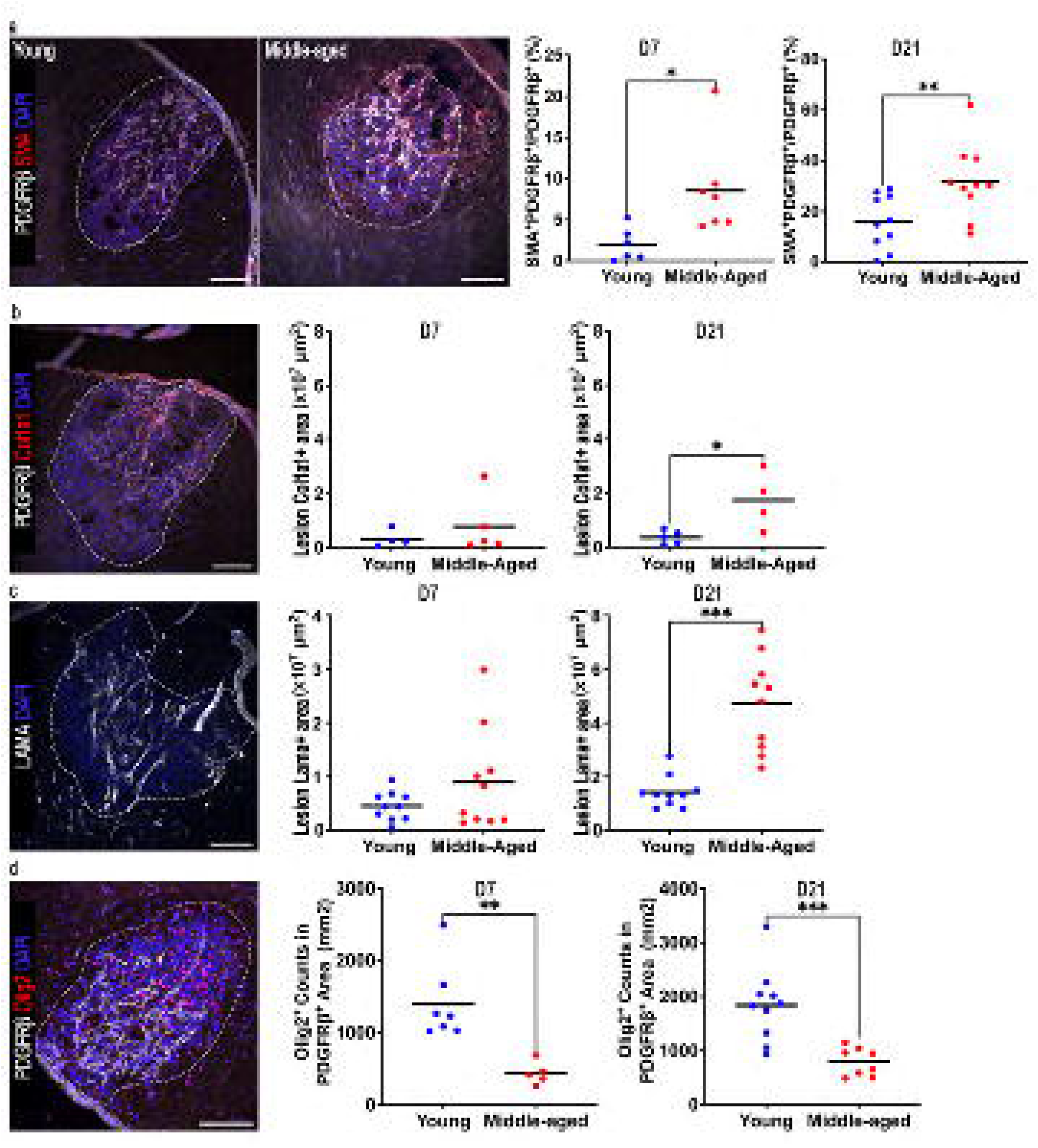
Age enhances fibroblast properties in demyelinated lesions. (**a)** Representative immunofluorescence images of day 7 LPC lesions in young and middle-aged mice labeled for DAPI, PDGFRβ and SMA; and quantification of the proportion of PDGFRβ overlapping with SMA at 7 and 21 dpl in young and middle-aged mice. (**b)** Representative immunofluorescence images of day 7 LPC lesion of young mice labeled for DAPI, PDGFRβ, and Col1a1; and quantification of Col1a1 positive area at 7 and 21 dpl in young and middle-aged mice. (**c)** Representative immunofluorescence images of day 7 LPC lesion in young mice labeled for DAPI and LAMA (laminin); and quantification of LAMA positive area at 7 and 21 dpl in young and middle-aged mice. (**d)** Representative immunofluorescence images of day 7 LPC lesion in young mice labeled for DAPI, Olig2, and PDGFRβ; graphs comparing the number of Olig2 positive cells within PDGFRβ positive regions of the LPC lesion at 7 and 21 days post LPC injury. In the graphs, each dot is an individual mouse. Significance is indicated as \**P*<0.05, \*\**P*<0.01, ***P<0.001; unpaired, *t*-test, comparing young and middle-aged groups. Data presented as mean.

### Fibroblasts are present in MS lesions

We extended from murine data into the human demyelinating condition, MS. Frozen brain sections from autopsied MS cases collected in Montreal were stained for PDGFRβ as well as CD45 to identify leukocytes, and DAPI for cell nuclei (Fig 8a). We observed that PDGFRβ+ cells appeared close to CD45+ leukocytes (Fig 8a).

**Fig. 8.**
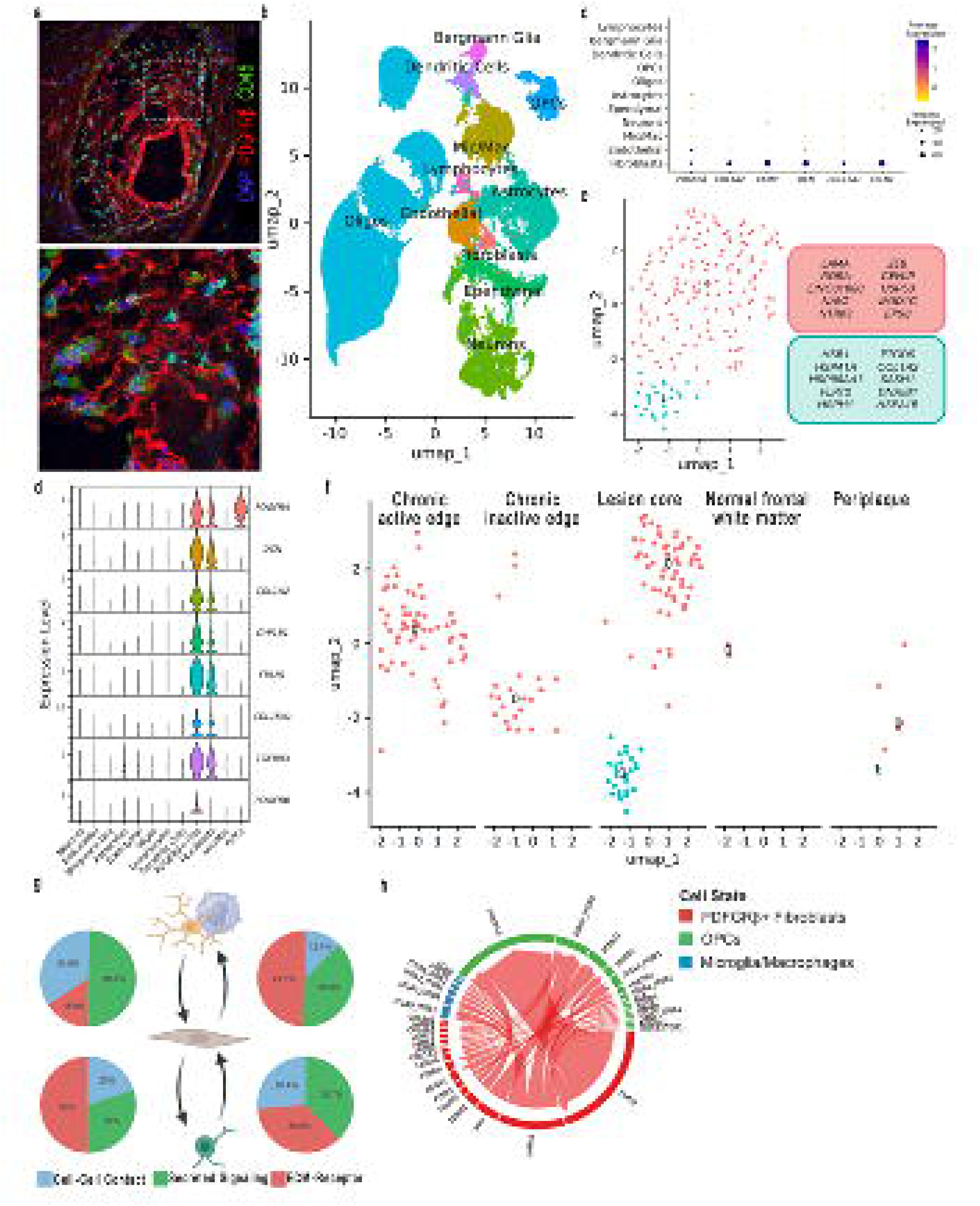
Fibroblasts in MS lesions. (**a)** Representative immunofluorescence images of MS lesions labeled for DAPI, CD45, and PDGFRβ. This is observed in 3 different MS cases from the Montreal Bank. (**b)** UMAP visualization of 66,432 cells from MS lesions and their cell annotation^58^. (**c)** Dot plot indicating markers used to identify fibroblast population. Size of the dot indicates the percentage of cells while the color encodes the average expression levels across all cells. (**d)** Violin plots depicting log normalized gene expression of fibroblast markers comparing PDGFRβ positive fibroblast population to other cell groups. (**e, f)**. UMAP visualization of (**e**) subclusters of PDGFRβ positive fibroblast population with top 10 DEGs, and (**f**) distribution of PDGFRβ positive fibroblast subclusters across MS lesion types and frontal white matter from neurologically normal controls. (**g**) Schematic indicating inferred outgoing and incoming fibroblast signals and their distribution across signaling databases. (**h**) Chord plot showing L-R interactions with fibroblasts as the source of signals.

Next, we reanalysed previously published snRNAseq datasets from human brain tissues of MS lesion rim, core and periplaque; and white matter tissue from neurologically healthy controls^58^. Unsupervised clustering identified 25 clusters from the 66,432 total cells including leukocytes, Bergmann glia, dendritic cells, vascular endothelial cells, OPCs, oligodendrocytes, astrocytes, ependymal cells, microglia/macrophages, and neurons (Fig 8b). We highlighted a cluster of 174 cells that expressed fibroblast related genes^59^ such as *PDGFRB, COL1A2, COL15A1,* and *DCN* (Fig 8c). From this cluster we focused on *PDGFRB+* cells as our population of interest that were positive for the markers *PDGFRA, COL1A2, TGFBR3, COL15A1*, and *LAMC3* (Fig 8d; Supplementary Fig 5a). Importantly, genes associated with pericytes (*RGS5*) and smooth muscle cells (*TAGLN*) were not appreciably expressed in our population of interest (Supplementary Fig 5b)^49^. The *PDGFRB+* fibroblasts were separated from the other cells in the dataset and re-clustered. This delineated 2 subpopulations (Fig 8e). Subcluster 1 expressed genes associated with stress response (*HSPB1, HMGB1*) while subcluster 0 expressed genes associated with ECM, cell proliferation, and migration (*LAMA2, NTRK3*) (Fig 8e). Within both clusters 0 and 1 are markers associated with perivascular (*CEMIP, RORA, NTRK3*) and meningeal fibroblasts (*COL1A2, PTGDS, FLRT2*)^53^. Assessment of these clusters via region showed that subcluster 0 was present in chronic active and chronic inactive edges and also the lesion core, but only minimally observed in control white matter or periplaque area; subcluster 1 was elevated only in the lesion core (Fig 8f).

### Fibroblast interactions in MS lesions

We interrogated the interactions of fibroblasts with other cell types in MS lesions using CellChat ligand-receptor interaction analysis. Similar to LPC lesions, fibroblasts, microglia/macrophages and OPCs all communicated with each other (Supplementary Fig 5c-f; Supplementary Table 4). Fibroblast outgoing signals primarily involved ECM-receptor interactions (Fig 8g) such as COL1A2-ITGAV/ITGB9 and LAMA2-CD44 (Fig 8h) while most incoming signals were via secreted signaling (Fig 8g; Supplementary Fig 5g). This is seen in the ligand-receptor pairings engaged by fibroblast derived ligands such as *COL1A2-ITGAV/ITGB9* and *LAMA2-CD44* (Fig 8g; Supplementary Table 4). As well, incoming secreted ligand-receptor interactions that fibroblasts were predicted to respond to included *FGF1-FGFR4* and *PDGFB-PDGFRA* (Supplementary Fig 5g; Supplementary Table 4).

Fibroblast ligand-receptor pairs were further categorized into signaling pathways such as transforming growth factor beta (TGFB), PDGF, FN1, and COLLAGEN (Supplementary Fig 5h, i). Fibroblasts contributed to pathways that drove microglia/macrophage signaling including COLLAGEN, LAMININ, CSF, and CXC motif chemokine ligand (CXCL) (Supplementary Fig 5h, i; Supplementary Table 4). Additionally, fibroblasts participated in pathways that signaled at OPCs including WNT, FGF, COLLAGEN, and LAMININ (Supplementary Fig 5h, i; Supplementary Table 4).

Comparing predicted fibroblast signaling across data sets highlights unique and conserved pathways (Supplementary Fig 6; Supplementary Table 5). When comparing MS to pathways shared by all the LPC datasets, we find only limited unique outgoing (HGF, MHC-I, ANNEXIN) and incoming (MHC-II, CypA, CD96) fibroblast pathways in MS (Supplementary Fig 6; Supplementary Table 5). Conversely, there were 55 shared outgoing and 56 shared incoming fibroblast signaling pathways between MS and the LPC datasets (Supplementary Fig 6). Shared outgoing pathways include those important for fibrosis and wound regeneration (COLLAGEN, FN1), OPC response (PDGF, PTN), and immune modulation (CSF, CXCL) (Supplementary Table 5). As such, shared fibroblast ligands predicted to act on OPCs are associated with diverse outcomes for OPC responses. Some ligands are inhibitory of OPC migration (*TNC, COL1A1*), proliferation (*HSPG2, FN1*), and maturation (*COL1A1, FGF2*) while some promote migration (*FN1, PDGF*), proliferation (*FGF2, BDNF*) and maturation (*LAMA2, PTN*) (Supplementary Table 5). Also, fewer interactions between microglia/macrophage and fibroblasts were predicted but include *IGF1-IGF1R, SPP1-ITGA5/ITGB1,* and *TGFA-EGFR/ERBB2* (Supplementary Table 5). Conversely, fibroblast response to microglia/macrophages ligands were associated with fibrotic wound responses (*TGFA-EGFR/ERBB2*^60^*, TGFB1-ACVR1/TGFBR1*), fibroblast proliferation (*PDGFA/B-PDGFRA/B*), acquisition of myofibroblast phenotype (*PDGFA/B-PDGFRA/B*^61^*, TGFB1-ACVR1/TGFBR1*), and stimulation (*LGALS9-CD44*^62^*, SPP1-ITGA5/ITGB1*^63^) (Supplementary Table 5). Interestingly, microglia/macrophage ligands *LGALS9* and *SPP1* interactions with fibroblast *CD44* are common only between MS and 10 dpl LPC, while *IGF1-IGF1R* and *PDGFA-PDGFRB* are shared between MS and 5 dpl LPC (Supplementary Table 5). This suggests that changes to microglia/macrophages phenotypes have potential consequences for altering fibroblast responses. Taken together these data highlight not only the presence of fibroblasts in MS lesions but also their prominent interactions with microglia/macrophages and OPCs in their environment.

### Assessment of fibroblasts in different lesion types of MS by immunofluorescence microscopy

Our previous immunofluorescence detection of PDGFRβ+ cells in MS specimens from the Montreal bank utilized only a single marker (PDGFRβ, Fig 8a), but this provided the rationale to interrogate transcriptomic databases that corroborated the presence of fibroblasts in MS lesions (Fig 8b-f; Supplementary Fig 5). To increase confidence that cells detected by immunofluorescence microscopy were fibroblasts, we subjected MS specimens from a different source, the Imperial College brain bank, to PDGFRβ+Col1a1+ double immunofluorescence analyses. Furthermore, different lesion types were examined. Figure 9 shows that PDGFRβ+Col1a1+ fibroblasts are found in active and chronic active lesions, in areas that are perivascular as well as sparsely distributed in the parenchyma. These results are observed in 5 of 5 active, and 5 of 5 chronic active lesions.

**Fig. 9.**
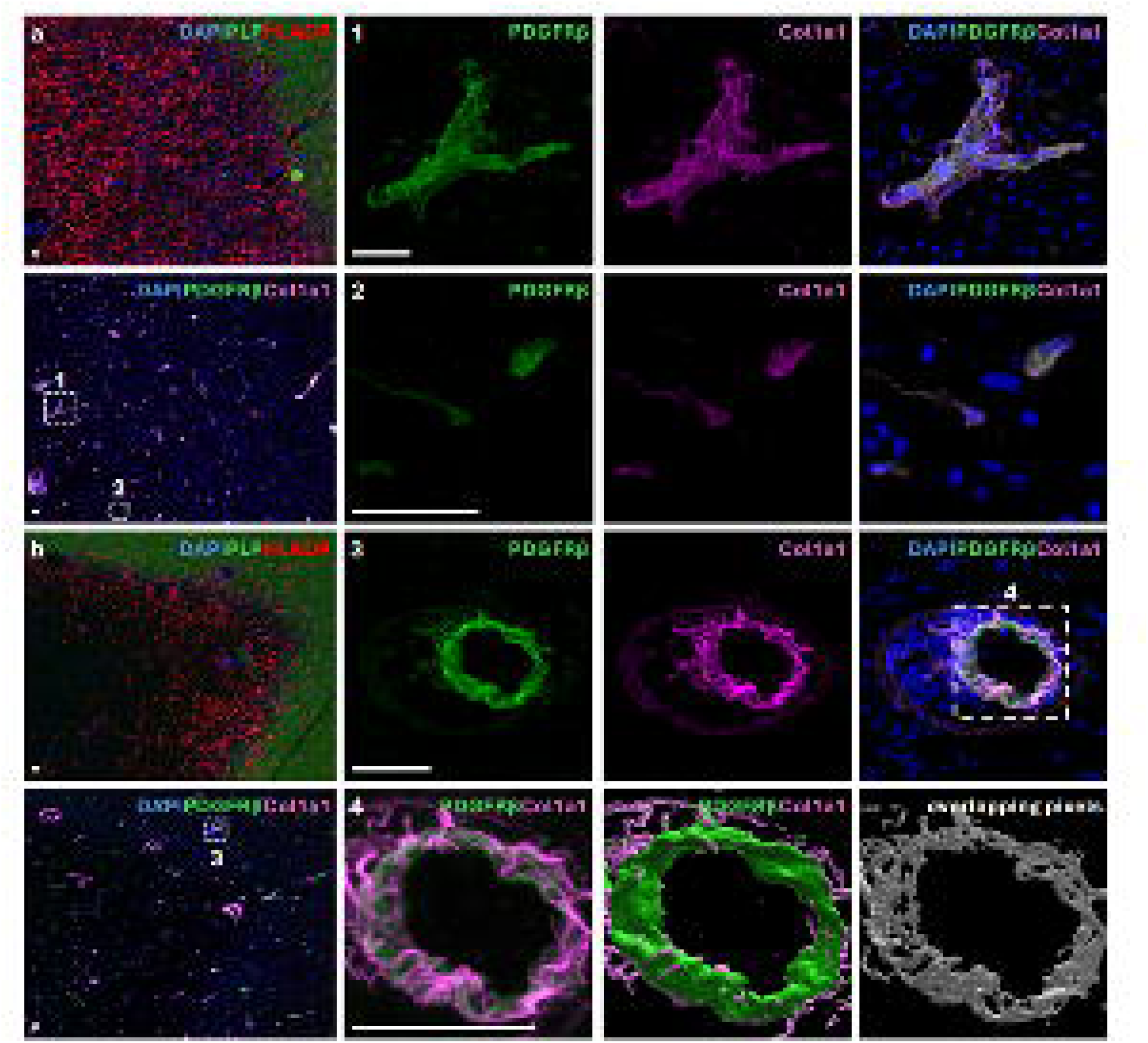
Fibroblasts are found in active and chronic active MS lesions. The nature of demyelinated (PLP-negative) lesions was identified by HLADR+ microglia/macrophages uniformly throughout a plaque (active) (**a**), or at the rim (chronic active) (**b**). Panels 1-3 show higher magnification inserts. PDGFRβ+Col1a1+ cells were detected in both lesion types, associated with vascular structures (e.g. 1 in a, 3 in b) or in parenchyma (e.g. 2 in a). These results were observed in 5 of 5 active, and 5 of 5 chronic active lesions. Panel 4 presents an Imaris 3D reconstruction demonstrating close spatial localization and partial overlap of PDGFRβ and Col1a1. Scale bars: 50μm.

## Discussion

The presence of parenchymal fibroblasts in both neuroinflammatory and traumatic CNS injury is more appreciable than previously thought^3,9^. Here, we report that the expansion of fibroblasts in the lysolecithin (LPC)-injured spinal cord white matter is correspondent with lowered oligodendrocyte density and exacerbated with age. We verify the identity of fibroblasts using PDGFRβ immunoreactivity and a PDGFRβ^TdTom^ reporter mouse line^45,46,64^, and by additional markers including fibronectin, Col1a1 and periostin. Moreover, transcriptomics data identified additional markers of fibroblasts, including several collagens, decorin and fibulin-1, and their absence of the pericyte marker Rgs5 and smooth muscle cell marker Talgn. Analysis of a snRNAseq dataset of LPC lesions shows that while fibroblast counts expand significantly at 5, 10, and 20 dpl, pericytes are scarce in the LPC lesion environment. Highly multiplexed RNA in situ hybridization (RNA-ISH, CosMx) further supports that fibroblasts are expanded in LPC lesions (16-fold increase) compared to sham controls.

In MS lesions, we identified PDGFRβ+Col1a1+ cells by immunohistochemistry in close proximity to CD45 positive leukocytes, in both active and chronic active lesions. We analysed a single-nucleus RNA sequencing dataset of MS^58^ and found a small population of presumptive fibroblasts. We identified a population of PDGFRβ positive cells that expressed genes associated with fibroblasts, and similar to those found in LPC lesions, such as *COL1A2, COL15A1, DCN* and *FBLN1*. It is noteworthy that these cells are found in lesions but minimally in the periplaque area or neurologically normal white matter, indicating their association with pathology in MS. This is consistent with findings from other groups that have shown greater presentation of PDGFRβ cells in chronic active MS lesions^14^. Cellchat analysis of both LPC and MS lesions indicates that fibroblasts possess immense potential to interact with a range of cells including microglia/macrophages and oligodendrocyte lineage cells in the lesion environments.

The potential effect of fibroblasts on CNS injuries has received scant attention, and little is known regarding their effect on the most common form of CNS regeneration, remyelination^3,6^. Others have suggested that fibroblasts have the potential to influence oligodendrocyte lineage cells that are responsible for remyelination^3,6^. Our analysis of predicted ligand-receptor interactions from CosMx suggests that fibroblasts are a potent driver of signaling in LPC demyelination including many fibroblast-derived ligands that can affect OPCs such as Fn1 and Col1a1^65^. Accumulation of inhibitory ECM components, such as fibronectin aggregates^21,66,67^, and collagen 1^68^ can impair OPC function, maturation, and reduce remyelination. ECM components can also impede OPC function through indirect effects including by stimulating microglia/macrophages and lymphocytes that kill OPCs^67^. Whether fibroblast phenotypes and their communication with other cells in LPC and MS lesions changes over time is unclear and requires further investigation. However, recent findings in traumatic brain injury have shown that fibroblasts do have dynamic responses over the development of the injury^9^. Furthermore, how potential fibroblast-led signaling interactions continue to influence remyelination requires further investigation, as fibroblasts are transcriptionally dynamic throughout remyelination^53^.

It is feasible that fibroblasts provide a physical obstacle to oligodendrocyte processes contacting axons for remyelination. Indeed, we show in vivo that few OPCs or newly formed myelin are present in areas occupied by fibroblasts despite persistence of axons. Our results in culture show that OPCs avoid fibroblast areas. Indeed, CellChat predicts fibroblast-to-OPC interactions to be amongst the most common. Nonetheless, it is difficult from our current data to infer mechanistically how fibroblasts affect OPCs, as the predicted fibroblast ligands that affect OPCs include factors inhibitory for OPC functions such as *Fn1* and *Col1a1*, but also others that support their activity including *Lama2* and *Ptn*. Future experiments are necessary to provide clarity to the mechanisms regulating the exclusion of oligodendrocyte lineage cells by fibroblasts. The lesion environment is pleiotropic with many factors of diverse functions and other variables such as mechano-transduction which can also affect oligodendrocyte properties. Although fibroblasts are not present in large numbers following injury, their broad surface area, potent capacity to exclude oligodendrocytes from lesions, and their many potential interactions with microglia/macrophages position them to be important regulators of pathophysiology and recovery.

Age contributes to overall morbidity and the reduced efficiency of many biological processes^55^ including the formation of oligodendrocytes^69^ and remyelination^22,70,71^. We found a greater number of fibroblasts in LPC lesions of middle-aged compared to young mice at both early and late timepoints with a greater accumulation of ECM at later timepoints. It has been well described that the rate of accumulation of microglia/macrophages is reduced within lesions of aging mice compared to young mice. We have previously shown that there is an expansion of microglia/macrophages in aging lesions that resembles pro-fibrotic scar-associated macrophages that express *Spp1*, *Trem2*, *Cd63*, and *Fabp5*^9,24,30,72,73^. Importantly, osteopontin (*Spp1*) stimulates inflammatory microglial states and exacerbates CNS injury, and Spp1-positive macrophages have been shown to drive the activation of myofibroblasts leading to increased tissue fibrosis^74^. Interestingly, this same population has been described in chronic CNS lesions including chronic active MS lesions^75^. This is consistent with our data in which we found that Spp1-positive microglia/macrophages are present in both our LPC dataset and the reanalyzed MS lesion results. Further interrogation of the connection between aging microglia phenotypes and fibrosis will provide more direct connection between this aging associated microglia/macrophage population and fibroblast expansion.

The consequences of the age associated expansion of fibroblasts are also of great interest. We found a reduction in the density of oligodendroglia in the fibroblast occupied areas. However, it has been known for some time that age impacts OPCs and remyelination efficiency. Without targeting the fibroblast response in the aging lesion, it is not certain that the reduced efficacy of repair in aging is a fibroblast driven phenomenon rather than an OPC or other age-related mechanisms such as immune responses. CNS lesions are known to become stiffer with age which contributes to OPC dysfunction^57^. We also found that the expansion of fibroblasts is associated with increased ECM levels. It is not clear if fibroblasts act directly or indirectly on OPCs in the aging lesion, but CellChat analysis suggests fibroblast-OPC interactions are primarily ECM mediated lending to a more indirect effect. Thus, the increased presence of fibroblasts in middle-aged LPC lesions may contribute to the worse remyelination outcomes noted with aging.

Fibroblasts undoubtedly expand in LPC lesions, however they remain a relatively rare cell type within the broader CNS, especially in homeostatic conditions. During homeostasis and injury fibroblasts are found surrounded by ECM which makes isolating them technically challenging. This is made more difficult with age as cells tend to be more sensitive to perturbation, making deeper investigation of age-related fibroblast changes difficult. Attempts to isolate and maintain meningeal fibroblast cultures were not successful limiting the study of aged fibroblast responses and interactions in vitro. This may be due to greater terminal differentiation of fibroblasts, loss of proliferative capacity, or increased sensitivity to environmental cues which have all been seen in other fibroblast populations in age^76,77^. By utilizing snRNAseq analysis of LPC allowed us to avoid some of these limitations. However, the LPC lesion environment is densely compacted with heterogeneous cell populations making individual cell segmentation difficult.

There are several limitations of our study. While fibroblasts expand in LPC lesions, they remain a relatively small cell type within the broader CNS, albeit with strong potential to influence their microenvironment. The scarcity of fibroblasts limits our capacity to subcategorize them into definitive subclusters, and to properly define the functions of each subtype; the low number of cells available for CellChat analyses in the spatial transcriptomics also reduces the reliability and stringency of the ligand-receptor pairing. Isolating fibroblasts for studies is also challenging. During homeostasis and injury, fibroblasts are surrounded by ECM which makes isolating them technically difficult, especially with age. Attempts to dissociate and maintain meningeal fibroblast cultures from aged brains were not successful, thus limiting the age-related characterization of fibroblast responses and mechanistic interactions in vitro. This may be due to greater terminal differentiation of fibroblasts, loss of proliferative capacity, or increased sensitivity to environmental cues which have all been seen in other fibroblast populations in age^76,77^. Another limitation is the curated 1000-plex panel of the CosMx spatial transcriptomics of LPC lesions, thereby allowing only a partial view of the properties of fibroblasts. Whole transcriptome single-cell resolution spatial transcriptomics with multiple samples will be needed in the future to fully examine fibroblast responses, including with aging, in situ.

In conclusion, we provide evidence that fibroblasts accumulate in demyelinated lesions including in MS. We show that fibroblast-occupied areas lack new myelinating cells, and that fibroblasts inhibit OPC differentiation in vitro. Importantly, this process is exacerbated by age leading to greater fibrosis of the lesion. These results highlight the necessity of targeting fibroblasts to mediate regenerative processes and improve outcomes in neurological conditions.

## Acknowledgements

We thank the Hotchkiss Brain Institute AMP core facility, and the NeuroOmics core facility, for technical support.

## Funding

We acknowledge operating grant support from MS Canada (number 1192262), the Canadian Institutes of Health Research (CIHR, FDN 167270) and National Natural Science Foundation of China (W2541024). BML and DM received PhD studentship awards, and RJ and RPG postdoctoral fellowships, from MS Canada. RPG also acknowledges the Dutch MS Society for the Gemmy and Mibeth Tichelaar Award. YD and SG were recipients of postdoctoral fellowships from CIHR. MM received an Internationalisation fellowship from the Carlsberg Foundation, Denmark.

## Competing interests

The authors report no competing interests.

## Data availability

The LPC single-nucleus RNA sequencing data are available from JKH upon reasonable request. All other data are available from VWY upon reasonable request. The mouse single-nucleus RNA sequencing datasets of LPC lesions are deposited in Gene Expression Omnibus (GEO) database under accession code GSE278643, and also bio-archived^53^. The CosMx data files can be accessed at the Figshare public database through accessing the link: https://figshare.com/s/61098937878dcfdb204f.

## Author contributions

BML produced the majority of the datasets and wrote the initial drafts of the manuscript. GSM, CDM, RPG, YD, SG, CL, MM, DM, RJ, PEM, CC Ling, CC-Lemarroy, RL and JKH provided data or experimental samples. VWY supervised and completed the final version of this manuscript. All authors read, edited and approved the final manuscript.

---

**Supplementary Figure 1.**
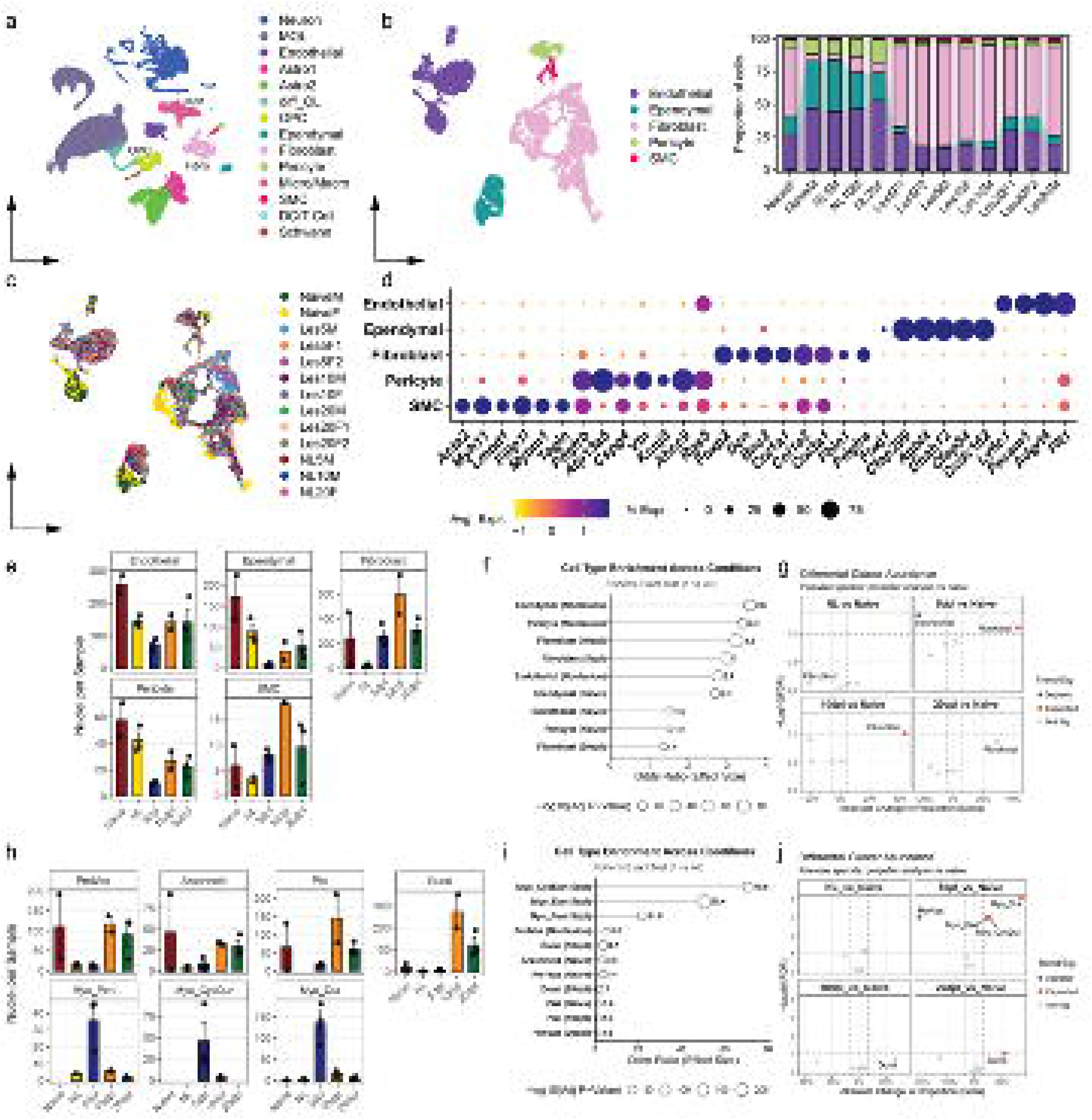

**Supplementary Figure 2.**
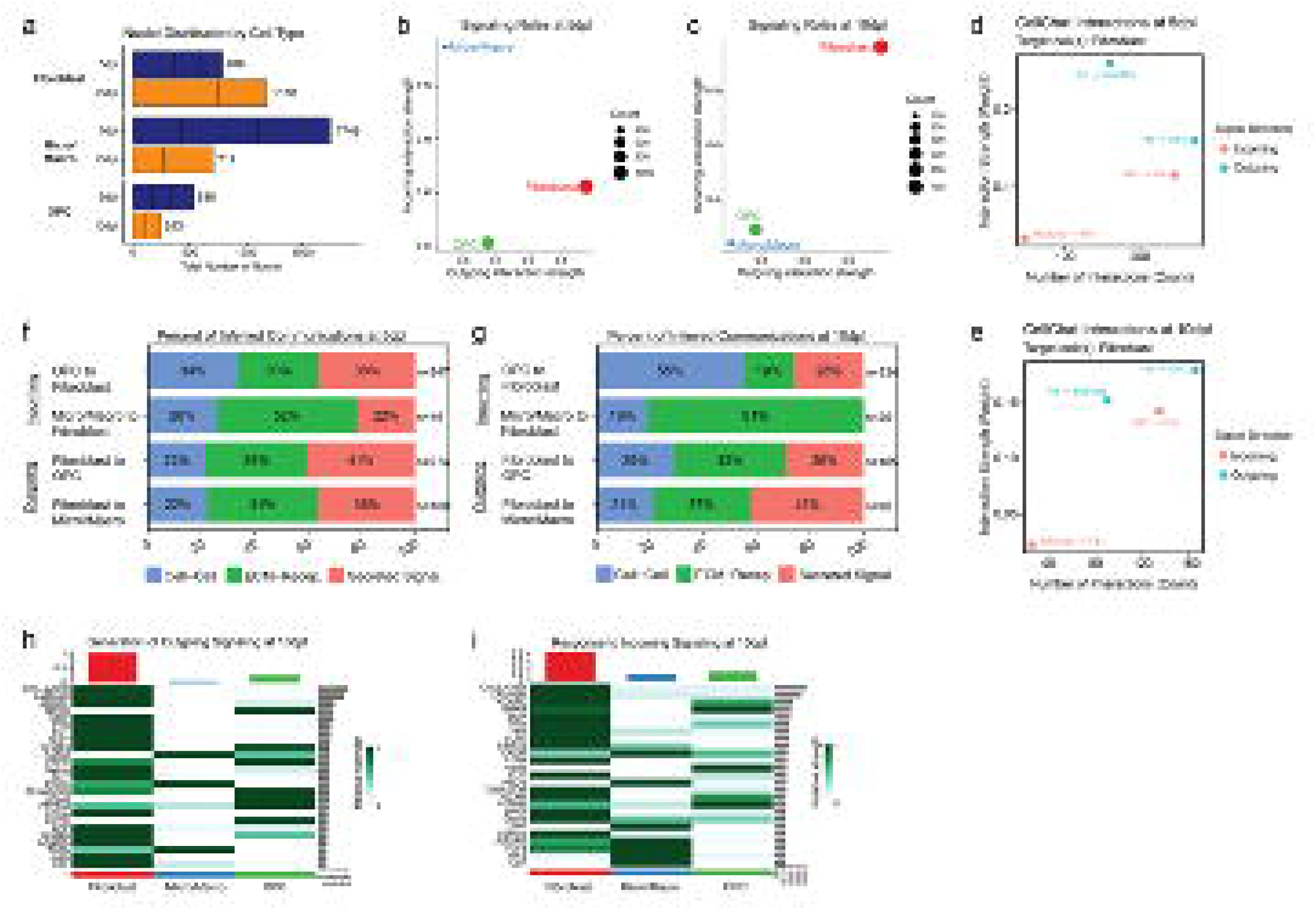

**Supplementary Figure 3.**
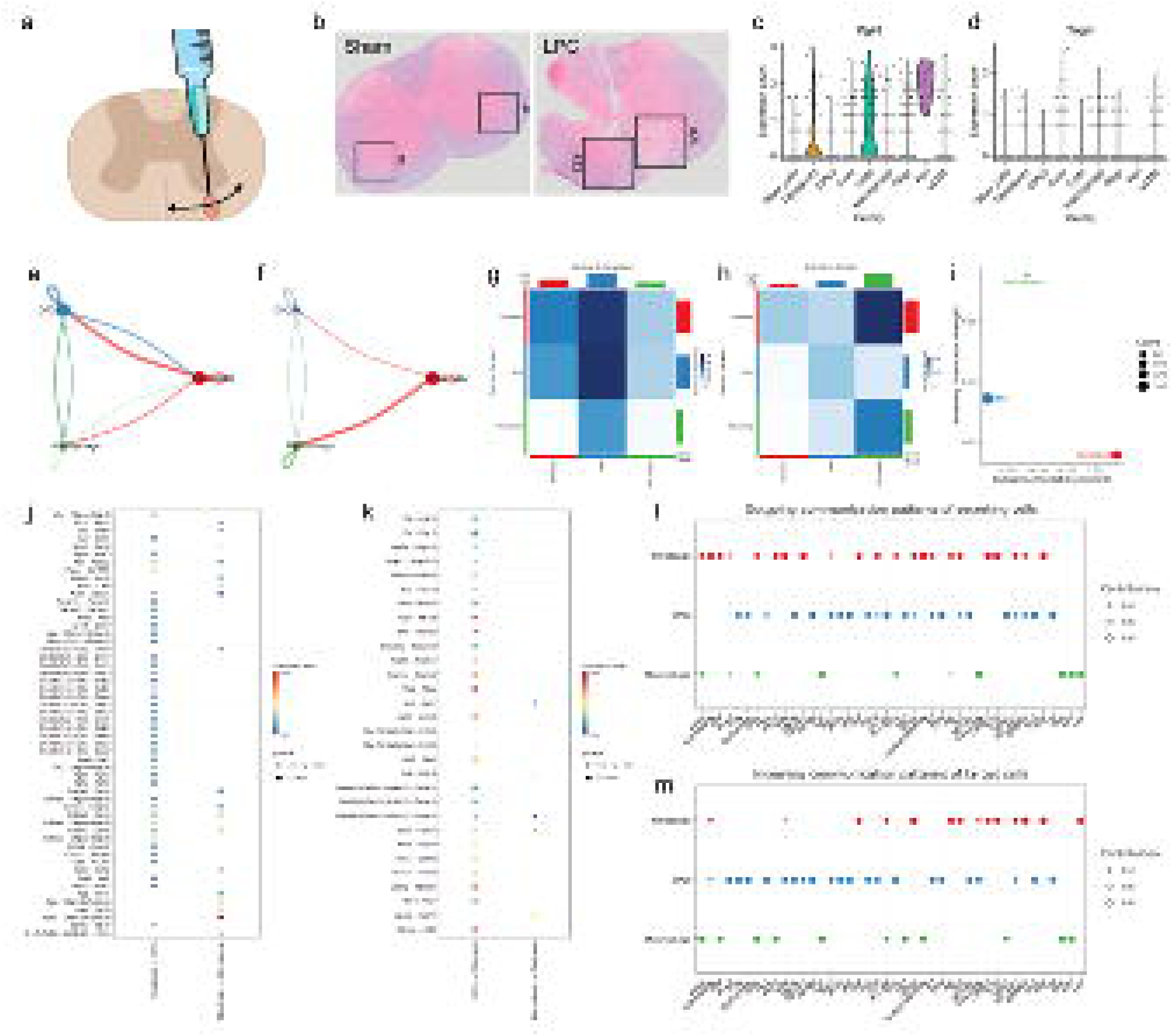

**Supplementary Figure 4.**
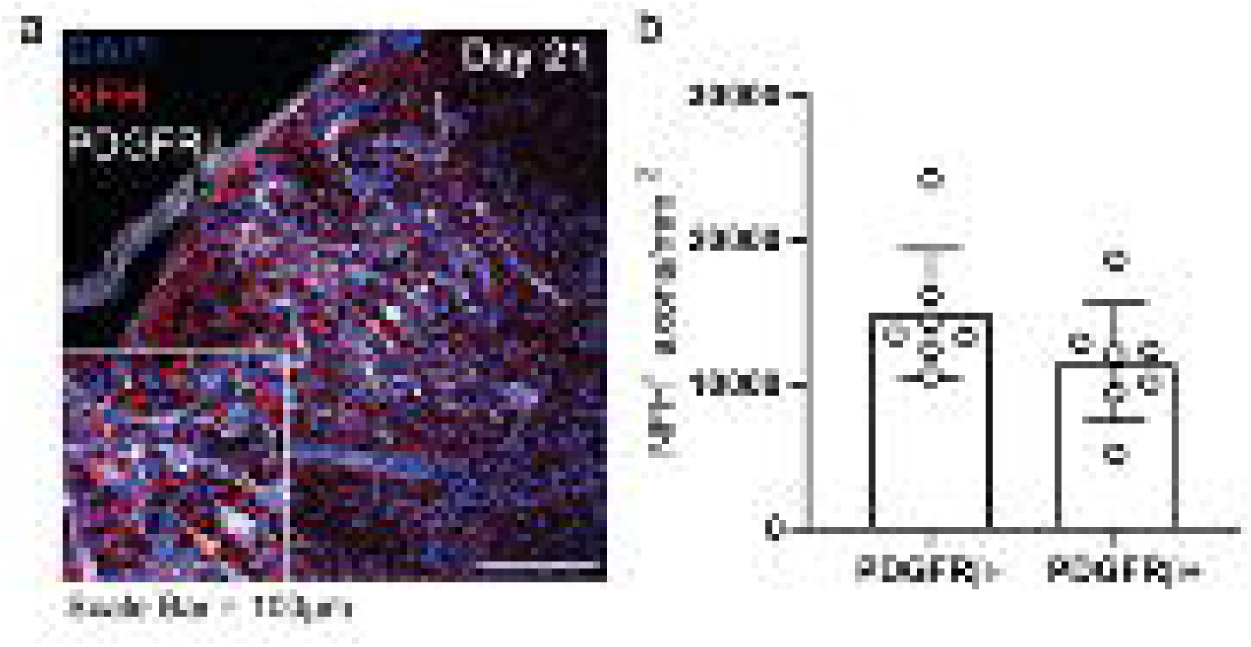

**Supplementary Figure 5.**
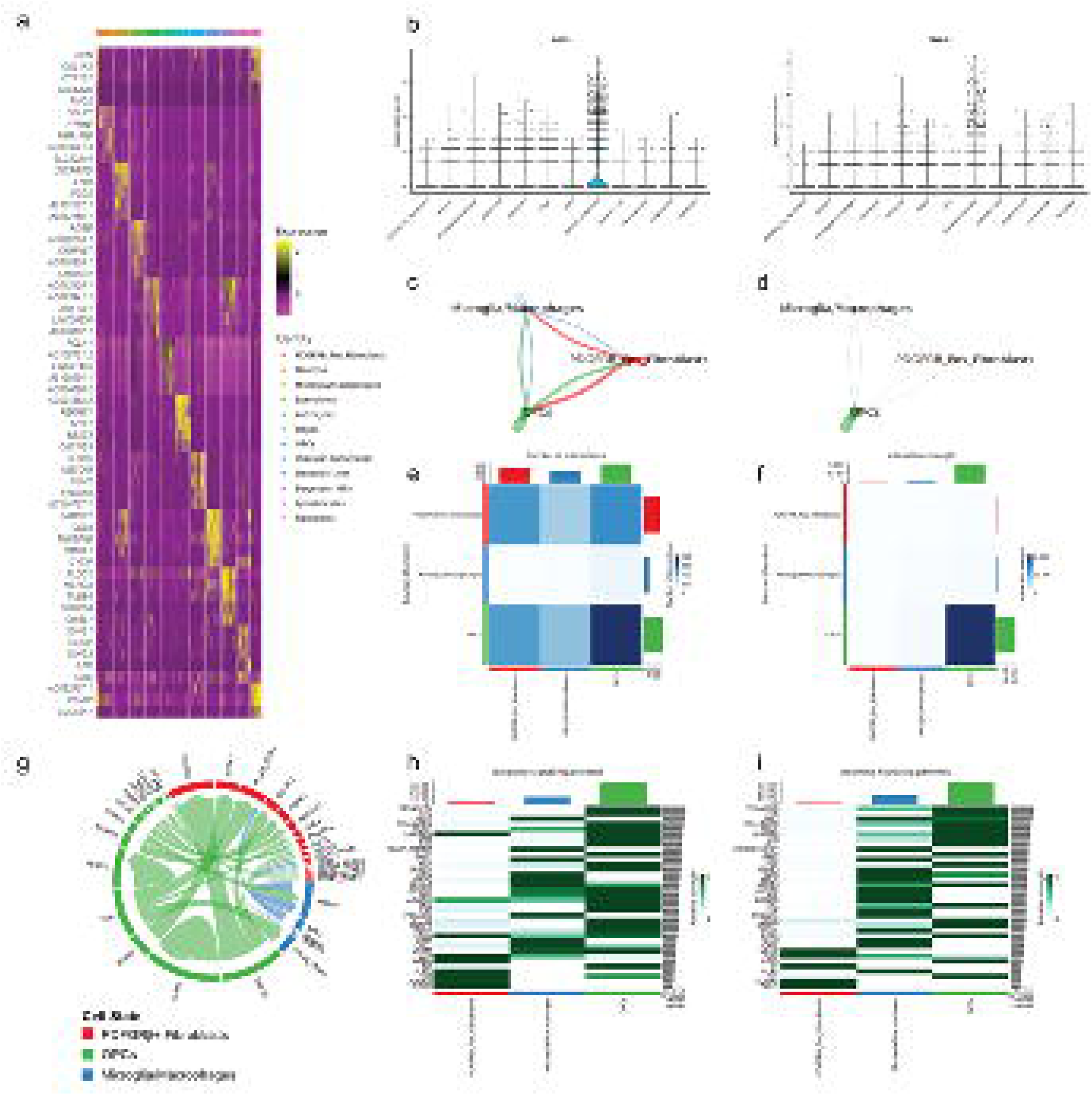

**Supplementary Figure 6.**
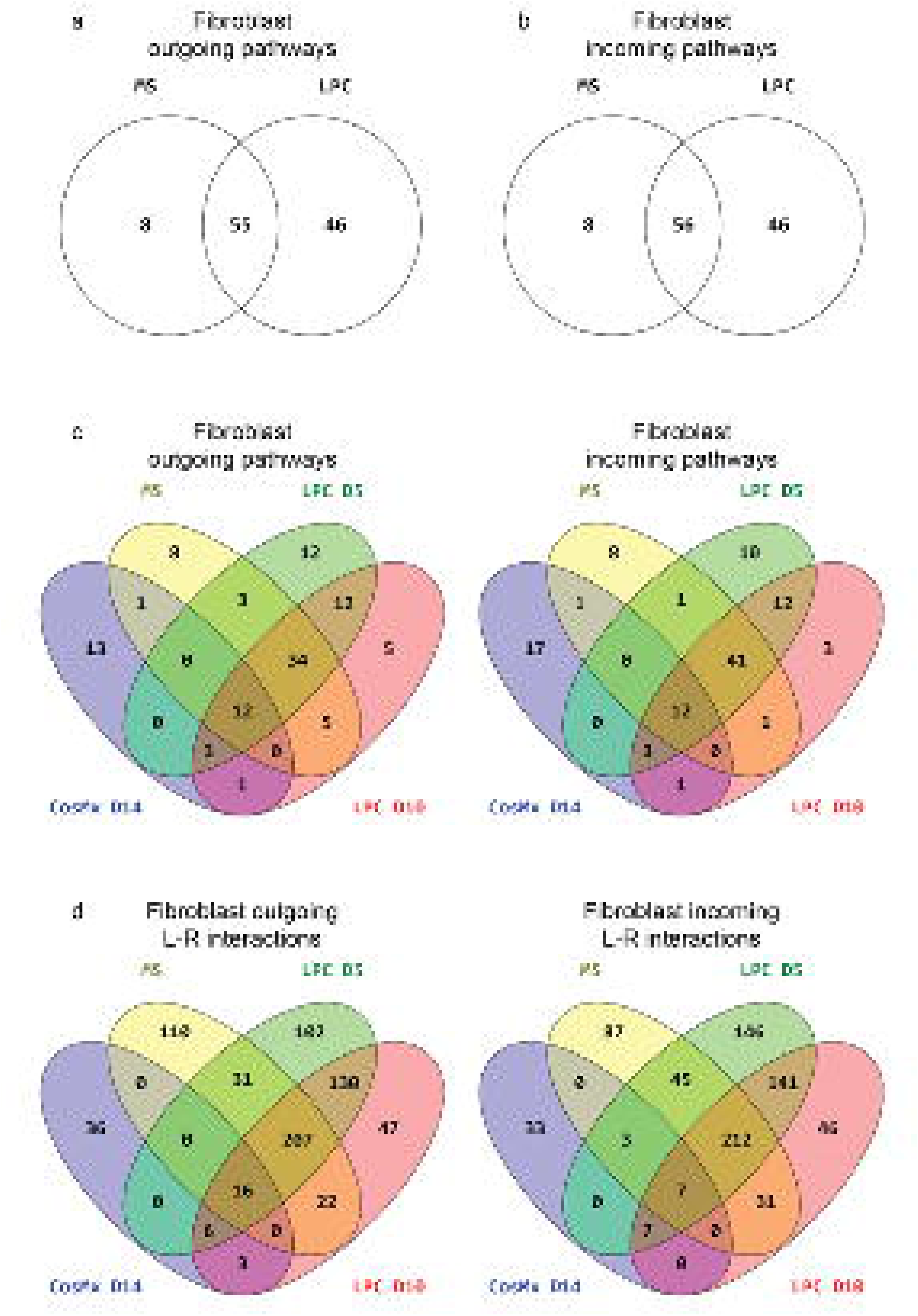

## Notes

### Competing Interest Statement

The authors have declared no competing interest.

### Summary of Updates

The manuscript has been thoroughly updated and new data has been integrated. Figures 2, 3, 5, 6, 8, 9, and supplementary figures 1, 2, 3, 5, and 6 have been updated. Author's have been updated.

